# BRCA1 and BRCA2 germline mutations driven signaling pathway alterations are sufficient to initiate breast tumorigenesis by the PIK3CA^H1047R^ oncogene

**DOI:** 10.1101/2023.07.26.550741

**Authors:** Poornima Bhat-Nakshatri, Aditi Khatpe, Duojiao Chen, Katie Batic, Henry Mang, Christopher Herodotou, Patrick C. McGuire, Xiaoling Xuei, Hongyu Gao, Yunlong Liu, George Sandusky, Anna Maria Storniolo, Harikrishna Nakshatri

## Abstract

Single cell transcriptomics studies have begun to identify breast epithelial cell and stromal cell specific transcriptome differences between BRCA1/2 mutation carriers and non-carriers. We generated a single cell transcriptome atlas of breast tissues from BRCA1, BRCA2 mutation carriers and compared this single cell atlas of mutation carriers with our previously described single cell breast atlas of healthy non-carriers. We observed that BRCA1 but not BRCA2 mutations altered the ratio between basal, luminal progenitor and mature luminal cells in breast tissues. A unique subcluster of cells within luminal progenitors is underrepresented in case of BRCA1 and BRCA2 mutation carriers compared to non-carriers. Both BRCA1 and BRCA2 mutations specifically altered transcriptomes in epithelial cells which are an integral part of NF-κB, LARP1, and MYC signaling. Reduction of MYC signaling and translational machinery in BRCA1/2 mutant epithelial cells is reminiscent of embryonic diapause-like adaptation that occurs in drug tolerant populations of cells. Signaling pathway alterations in epithelial cells unique to BRCA1 mutations included STAT3, BRD4, SMARCA4, HIF2A/EPAS1, and Inhibin-A signaling. BRCA2 mutations were associated with upregulation of IL-6, PDK1, FOXO3, and TNFSF11 signaling. These signaling pathway alterations are sufficient to alter sensitivity of BRCA1/BRCA2-mutant breast epithelial cells to transformation as epithelial cells from BRCA1 mutation carriers overexpressing hTERT + PIK3CA^H1047R^ generated adenocarcinomas, whereas similarly modified mutant BRCA2 cells generated basal carcinomas in NSG mice. Thus, our studies provide a high-resolution transcriptome atlas of breast epithelial cells of BRCA1 and BRCA2 mutation carriers and reveal their susceptibility to PIK3CA mutation-driven transformation.

**Statement of Significance:** This study provides a single cell atlas of breast tissues of BRCA1/2 mutation carriers and demonstrates that aberrant signaling due to BRCA1/2 mutations sufficient to initiate breast cancer by mutant PIK3CA.

## Introduction

BRCA1 and BRCA2 are well characterized breast cancer susceptibility genes and it is well established that mutations in these genes impair the homologous recombination mediated DNA repair pathway, which is required to maintain genomic integrity (1). Additionally, it has been suggested that breast epithelial cells from BRCA1 and BRCA2 mutation carriers undergo accelerated aging (2). Breast epithelial cells from BRCA2 mutation carriers have also been shown to be susceptible to aneuploidy due to DNA damage with attenuated replication checkpoint and apoptotic responses and age-associated expansion of luminal progenitor compartment (3). Similar expansion of luminal progenitor cells in BRCA1-mutation carriers have also been reported previously (4,5).

Single cell DNA/RNA sequencing is now used to determine whether inherited mutations affect mutation frequency, cell composition, and differentiation trajectory in adult organs. For example, single cell DNA sequencing of telomerase immortalized human mammary epithelial cells with and without manipulation of the endogenous BRCA1/2 locus, as well as breast tissues from BRCA1/2 mutation carriers, has revealed a high frequency of single nucleotide variations and small deletions and insertions in BRCA1/2 mutation carriers compared to non-carriers (6). A single cell RNA sequencing (scRNA-seq) study involving tumor adjacent normal or prophylactic surgery of BRCA1 mutation carriers and three non-carriers suggested that breast cancers in BRCA1 mutation carriers originate from luminal progenitors, as suggested previously using flow cytometry and bulk RNA sequencing (4,7). In mouse models, BRCA1 deficiency has been shown to cause aberrant differentiation of luminal progenitors (8).

Most single cell RNA-sequencing studies of human breast tissues utilized tissues from reduction mammoplasty and/or normal adjacent to tumors as “normal” controls, which we and others have shown to be histologically abnormal with changes in cell composition and gene expression (9,10). For example, 71% of reduction mammoplasty samples demonstrate non-proliferative disease compared to 31% of breast tissues from clinically healthy women. We demonstrated that tumor adjacent normal breast tissues of women of European ancestry contain elevated numbers of ZEB1+ stromal cells, while these cells are intrinsically elevated in the breast tissues of women of African ancestry (10). We also demonstrated distinct gene expression differences between healthy normal, tumor adjacent normal, and tumor tissues (10). Similarly, others have demonstrated changes in DNA methylation and gene expression in tumor adjacent normal compared to healthy breast tissues (11). These comparative studies between breast tissues of clinically normal donors, reduction mammoplasty samples and tumor adjacent normal were possible due to the availability of the Komen Normal Tissue Bank (KTB), an institutional biorepository to which healthy women donate breast biopsies for research purposes. Using these tissues, we have created various tools for breast cancer research including multiple immortalized cell lines with luminal enriched gene expression patterns, a minimum requirement for transformation of these immortalized cell lines, and the generation of a single cell transcriptome atlas of the breast tissues from women without BRCA1/2 mutations (referred as non-carriers hereafter) (12–14). This single cell breast atlas of the non-mutation carriers allowed us to perform comparative analysis of breast tissues from BRCA1 and BRCA2 mutation carriers with that of breast tissues from non-carriers. Similar to data obtained in mouse models of BRCA1 deficiency (15), we observed constitutive activation of NF-κB signaling in both BRCA1 and BRCA2 mutated cells. Moreover, while the transformation of immortalized cells from non-carriers required a combination of mutant PIK3CA (PIK3CA^H1047R^) and SV40-T/t antigens (13), PIK3CA^H1047R^ alone was sufficient to transform immortalized cells from BRCA1 and BRCA2 mutation carriers. These results suggest that basal signaling pathway alterations due to BRCA1 or BRCA2 mutations reduce the threshold of other genomic aberrations required to initiate breast tumorigenesis.

## Materials and Methods

### Tissue samples for single cell analysis

All tissues for the study were obtained after informed consent and approval from the Institutional Review Board. The majority of tissues were obtained from women undergoing prophylactic mastectomy after curative surgery ± chemotherapy and histopathology did not detect any abnormalities. BRCA mutation status was extracted from clinical reports but data on specific point mutations are not available to investigators. Breast tissues were cryopreserved as described previously (16) and thawed just before single cell generation for sequencing or cell line generation. Additional details of breast tissues are provided in supplementary Table S1.

### Tissue Dissociation, cDNA library preparation, and sequencing

Sample preparation, dissociation, and single cell RNA sequencing of individual samples were performed as described previously (14). Although breast tissues from 13 BRCA1 and 9 BRCA2 carriers were subjected to single cell RNA sequencing, good quality data were obtained only from five samples from BRCA1 and four samples from BRCA2 mutation carriers. Within five BRCA1 carrier samples, two of them were from the same donor but from randomly selected regions of left and right breasts sequenced separately to determine if there is breast region-specific variation in single cell profiles. Furthermore, in the integrated data analysis, data from one BRCA1 sample highlighted in our previous study were included (14). Sequence alignment, individual and integrated data analyses have been described previously and utilized 10X genomics Loupe Browser (14). Integration of two datasets utilized FindIntegrationAnchors function. Dataset that compares scRNA-seq data of breast tissues of non-carriers with BRCA1 or BRCA2 mutation carriers can be visualized through the following link and differences in expression levels of individual genes can be verified using this link (http://clark.ccbb.iupui.edu//Hari_BRCA). Genes differentially expressed in various cell types between BRCA1 mutation, BRCA2 mutation and non-carriers were subjected to Ingenuity Pathway Analysis (IPA) to identify signaling networks specifically active in BRCA1-mutated and BRCA2-mutated cells. Single cell RNAseq data have been deposited in NCBI GEO with accession number GSE223886.

### Establishment of breast epithelial cell lines from BRCA1 and BRCA2 mutation carriers, oncogene overexpression and animal studies

An hTERT immortalized cell line from BRCA1 mutation carrier was established from benign breast tissue of a 35-years old White woman with no prior treatment, whereas the cell line from a BRCA2 mutation carrier was established from normal breast tissues of a 32-years old White woman with no prior treatment using previously established protocols (12). Immortalized cell lines were infected with specific oncogene expressing lentiviruses as described previously (13).

Indiana University Animal Care and Use Committee has approved all animal studies and all studies were conducted as per NIH guidelines. Five million cells in 50% Matrigel (Corning, 354234) in 100 µl volume were injected into the mammary fat pad of NSG mice. Mice were implanted with 60-day slow release estradiol (SE-121, 0.72 mg pellet, Innovative Research of America). Animals were monitored for up to three months for tumor formation. At the end of the study, tumors and lungs were collected and subjected to H&E and immunohistochemistry (IHC) as described previously (13). Antibodies used for IHC have also been described previously (13).

### Western blotting

Western blotting for PIK3CA, phospho-p65 and p65 using cell lysates prepared in RIPA buffer was done as described previously. PIK3CA antibody that preferentially recognizes H1047R mutant was purchased from Assay Biotechnology (Catalogue # V0111). Phospho-p65 antibody (Ser 536, Catalogue # mAB #3033) and p65 antibody (#3034) were purchased from Cell Signaling Technology. Blots were reprobed with an antibody against βActin (#A5441, Sigma-Alderich). While cells for PIK3CA detection were grown under regular growth media, pp65 and p65 were measured after serum starving cells overnight to reduce the influence of growth factors in the media on NF-κB activation. For unknown reasons, serum starvation caused robust induction of endogenous PIK3CA and differences in expression between vector control and PIK3CA overexpressing cells could not be measured.

### Quantitative Reverse Transcription Polymerase Chain Reaction (qRT-PCR)

Total RNA was extracted using RNAeasy kit from Qiagen (#74104) and 1 ug of RNA was used to synthesize cDNA using iScript cDNA Synthesis Kit (#1708891) from BioRAD. qRT-PCR was performed using TaqMan universal PCR mix (#4324018) and predesigned TaqMan assay primers (Applied Biosystems). The following primers were used. MIR205HG (Hs03405498) and ACTB (Hs01060665_g1)

### NF-κB inhibitor sensitivity assay

2000 cells/well were plated in 96 well plate. Cells were treated with indicated concentrations of dimethylaminoparthenolide (DMAPT) for 48 hours (17). The effect of DMAPT on cell proliferation was measured using the Bromodeoxyuridine (BrDU) incorporation-ELISA (Millipore, Cat. No. 2752) as per the manufacturer’s instructions.

### Statistical analysis

Graphpad Prism software was used for statistical analysis of tumor incidence and for statistical analysis of *in vitro* data.

## Results

### Generation of single cell atlas of breast tissues of BRCA1 and BRCA2 mutation carriers

We analyzed scRNA-seq data at the individual donor level as well as integrating data from all samples together. Representative data from several donors of BRCA1 and BRCA2 mutation carriers and integrated data are shown in Figure 1. In one BRCA1 mutation carrier case, data from the left and the right breast are shown to demonstrate similar cell composition in both breasts. Tissue was sampled from random regions of the breast and both showed similar cell composition. Similar to breast tissue of non-carriers (14), breast tissue of BRCA1/2 mutation carriers contained distinct populations of epithelial cells, endothelial cells, adipocytes, fibroblasts and multiple immune cell types. Number and percentage of each cell types in BRCA1 and BRCA2 mutation carriers compared to non-carriers are shown in Table S2. We noted a higher percentage of adipocytes in breast tissues of BRCA2 mutation carriers than others, although significance of this difference is unknown.

**Figure 1:**
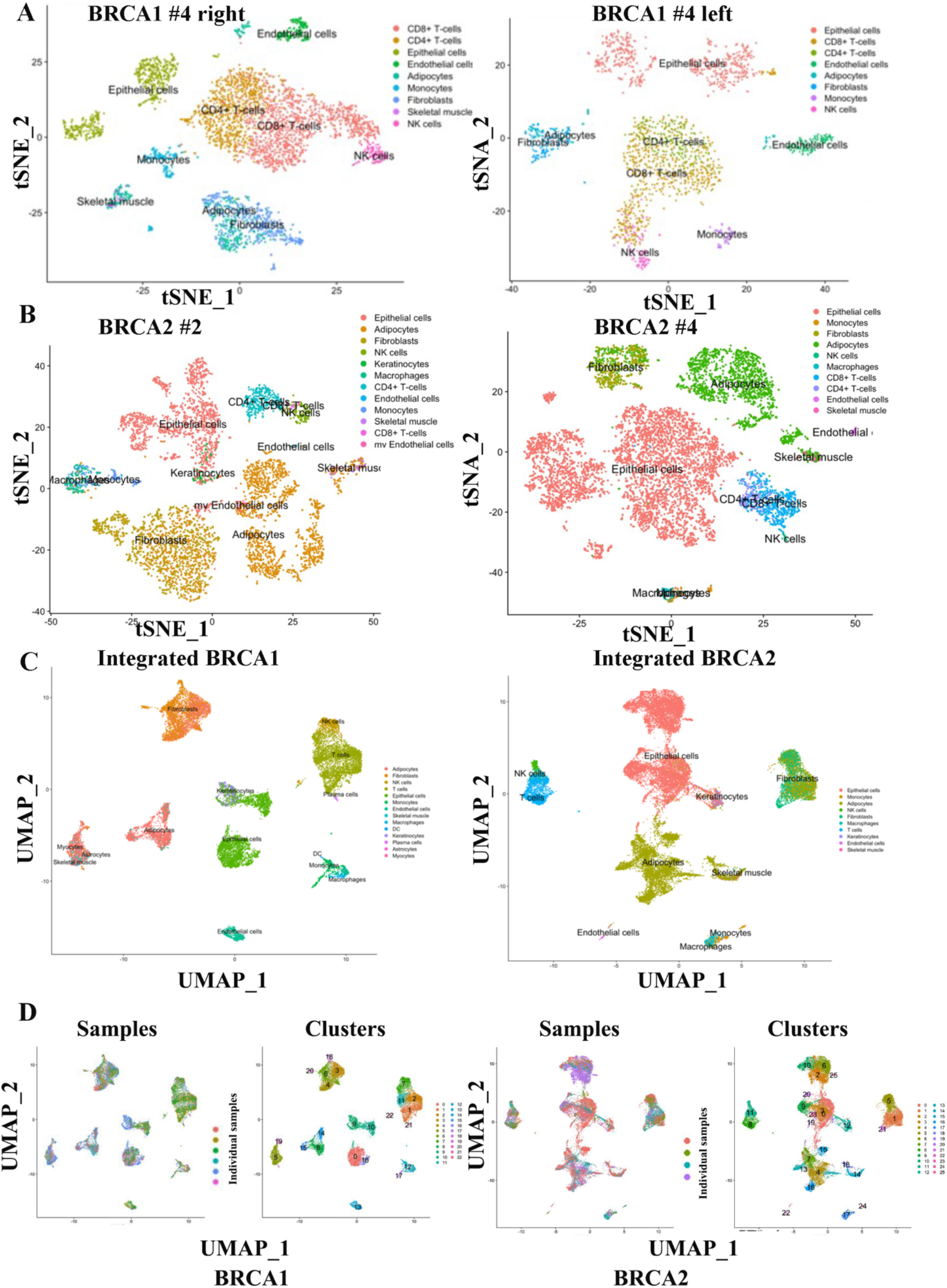
Single cell atlas of breast tissues of BRCA1 and BRCA2 mutation carriers compared to breast tissues of non-carriers. A) tSNE plots showing single cell map of the breast tissues from the right and left breast tissues of a 45 years old BRCA1 mutation carrier. B) tSNE plots showing single cell map of the breast tissues from 67 years old (left) and 39 years old (right) BRCA2 mutation carriers. C) UMAP showing integrated data from all samples sequenced. D) UMAP of individual sample overlay with cluster numbers.

We next overlayed scRNA-seq data of BRCA1 and BRCA2 mutation carriers with scRNA-seq data from breast tissues of non-carriers to determine whether there are any detectable differences in cell composition. Missing minor epithelial cell clusters were noted in the BRCA1 and BRCA2 mutation carriers compared to epithelial cell clusters generated from non-carriers (Figure 2). Between BRCA1 and BRCA2, one minor epithelial cell cluster (cluster 19) was missing in BRCA1 compared to BRCA2.

**Figure 2:**
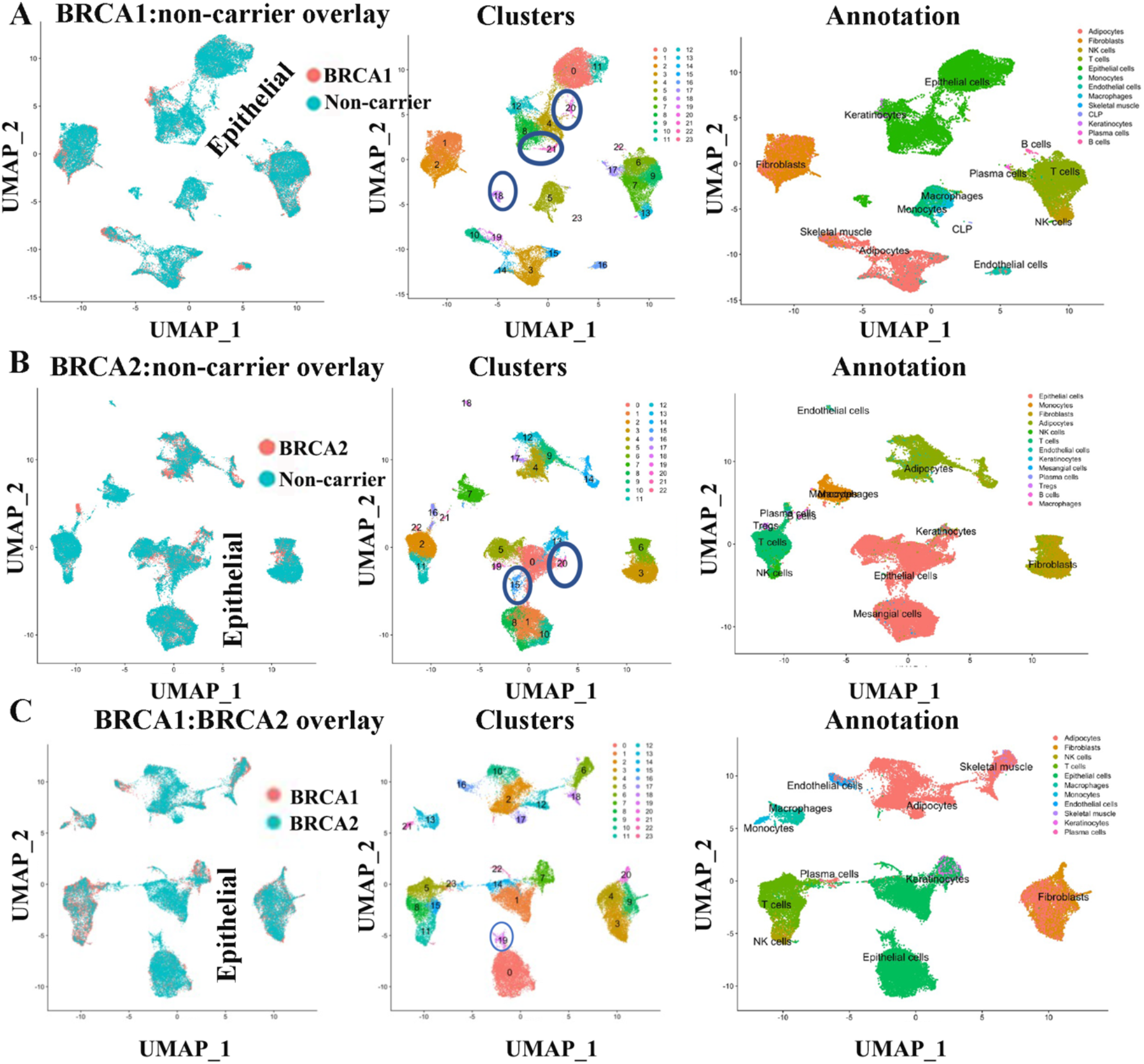
Overlay of BRCA1 and BRCA2 single cell map over the single cell map of breast tissues from non-carriers. A) Overlay of BRCA1 carrier-derived single cell atlas with that of non-carrier derived atlas. Non-carrier derived atlas has been described previously (14). Few minor clusters found in non-carrier atlas show limited overlap with BRCA1 mutation carrier-derived clusters (clusters 18, 20 and 21, oval in the center). All three are epithelial cell clusters. B) Overlay of BRCA2-derived single cell atlas with non-carrier derived atlas. Clusters 15 and 20, both epithelial clusters, are underrepresented in BRCA2 carriers. C) Overlay of BRCA1 carrier derived single cell atlas with that of BRCA2 carrier-derived single cell atlas. A minor cluster (cluster 19) is underrepresented in BRCA1 compared to BRCA2 carrier-derived single cell atlas.

### Breast tissues of BRCA1 and BRCA2 mutation carriers show differences in a subpopulation of luminal progenitors compared to breast tissues of non-carriers

Using the previously described markers of stem/basal (CD49f^+^/EpCAM^-^), luminal progenitors (CD49f^+^/EpCAM^+^) and mature luminal cells (CD49f^-^/EpCAM^+^) (18), we subclassified epithelial cells and compared these cells between three groups (Figure 3A-C). Two major differences can be seen. Despite a lower number of mature luminal and luminal progenitor cells, the basal/stem cell population in BRCA1 mutation carriers was higher compared to BRCA2 or non-mutation carriers (Figure 3D). While the percentage of basal cells in non-carrier and BRCA2 mutation carriers was 5.8% and 5.7%, respectively, it was 14.3% in the case of the BRCA1 mutation carriers. An increase in basal cells in BRCA1 mutation carriers was also reported in another recent study (7). We, however, did not observe significant differences in luminal progenitor cells between groups. Second, closely related luminal progenitor subclusters 10, 16 and 17 were missing in BRCA1 mutation carriers compared to non-carriers. Similarly, cluster 9 of the luminal progenitor is missing in BRCA2 mutation carriers. Clusters that are missing in BRCA1 and BRCA2 mutation carriers displayed higher expression of alveolar cell marker genes such as FOLR1 (19). These clusters in non-carriers also express the highest level of Osteopontin (also called SPP1). The top 10 genes that are differentially expressed in these missing clusters compared to other epithelial clusters of BRCA1and BRCA2 carriers and non-carriers are shown in Figure 3E. Except for differences in number of basal cells, there were no other major differences between epithelial cells of BRCA1 and BRCA2 mutation carriers.

**Figure 3:**
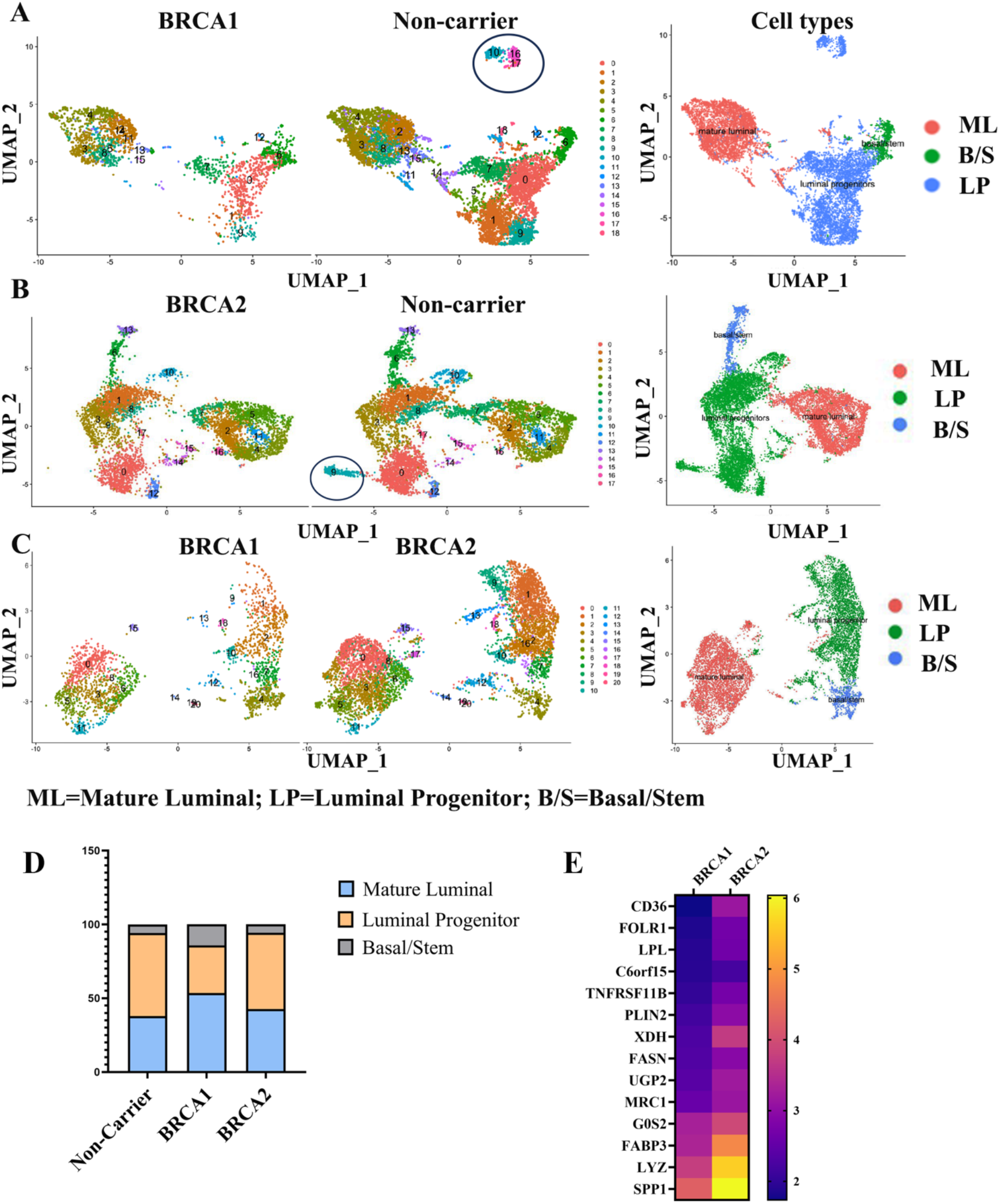
Comparison of breast epithelial cells of BRCA1 and BRCA2 mutation carriers with that of non-carriers. Breast epithelial cells were subclassified into basal/stem, luminal progenitor and mature luminal cells based on CD49f and EpCAM expression status as described previously (14). A) Side-by-side comparison of breast epithelial cells of BRCA1 mutation carriers with that of non-carriers. A distinct cluster of luminal progenitors is present only in non-carriers (indicated by an oval shape). B) Side-by-side comparison of breast epithelial cells of BRCA2 with that of non-carriers. Similar to BRCA1, a distinct cluster of luminal progenitor cells is present only in non-carriers. C) Side-by-side comparison of breast epithelial cells from BRCA1 mutation carriers with that of BRCA2 mutation carriers. D) Distribution pattern of basal/stem, luminal progenitor and mature luminal cells in non-carrier, BRCA1 and BRCA2 mutation carriers. E) Heatmap showing top 10 genes highly expressed in cluster missing in BRCA1 and BRCA2 mutation carriers compared to other clusters.

### Individual level gene expression differences between breast tissues of non-carriers and BRCA1/2 mutation carriers

Because BRCA1 and BRCA2 have transcription regulatory function through resolution of R-loops at transcription start sites (20), we next asked whether BRCA1 and BRCA2 mutations affect levels of individual transcripts. Towards this goal, we compared gene expression in epithelial, endothelial, and fibroblast cells of mutation carriers with non-carriers. All three cell types showed significant differences in expression of ∼100 genes (Table S3-S5). Differences between cells of BRCA1 and BRCA2 mutation carriers were minor (Table S5). Consistent with previous reports (7,21), epithelial cells of BRCA1 mutation carriers expressed higher levels of KRT14 compared to non-carriers (Table S3). Expression level differences in several genes, particularly in epithelial cells, are shown in Figure 4 and Figure S1. For example, CXCL13 is expressed at higher levels in a subpopulation of epithelial cells of non-carriers and BRCA2 mutation carriers but not in BRCA1 mutation carriers. MIR205HG is expressed at higher levels in epithelial cells of BRCA1 and BRCA2 mutation carriers but not in non-carriers. SERPINA3 is expressed at higher level in epithelial cells of BRCA1 and BRCA2 mutation carriers compared to non-carrier (Figure S1); this has previously been shown to confer invasiveness and epithelial to mesenchymal transition (EMT) phenotype to breast cancer cells (22).

**Figure 4:**
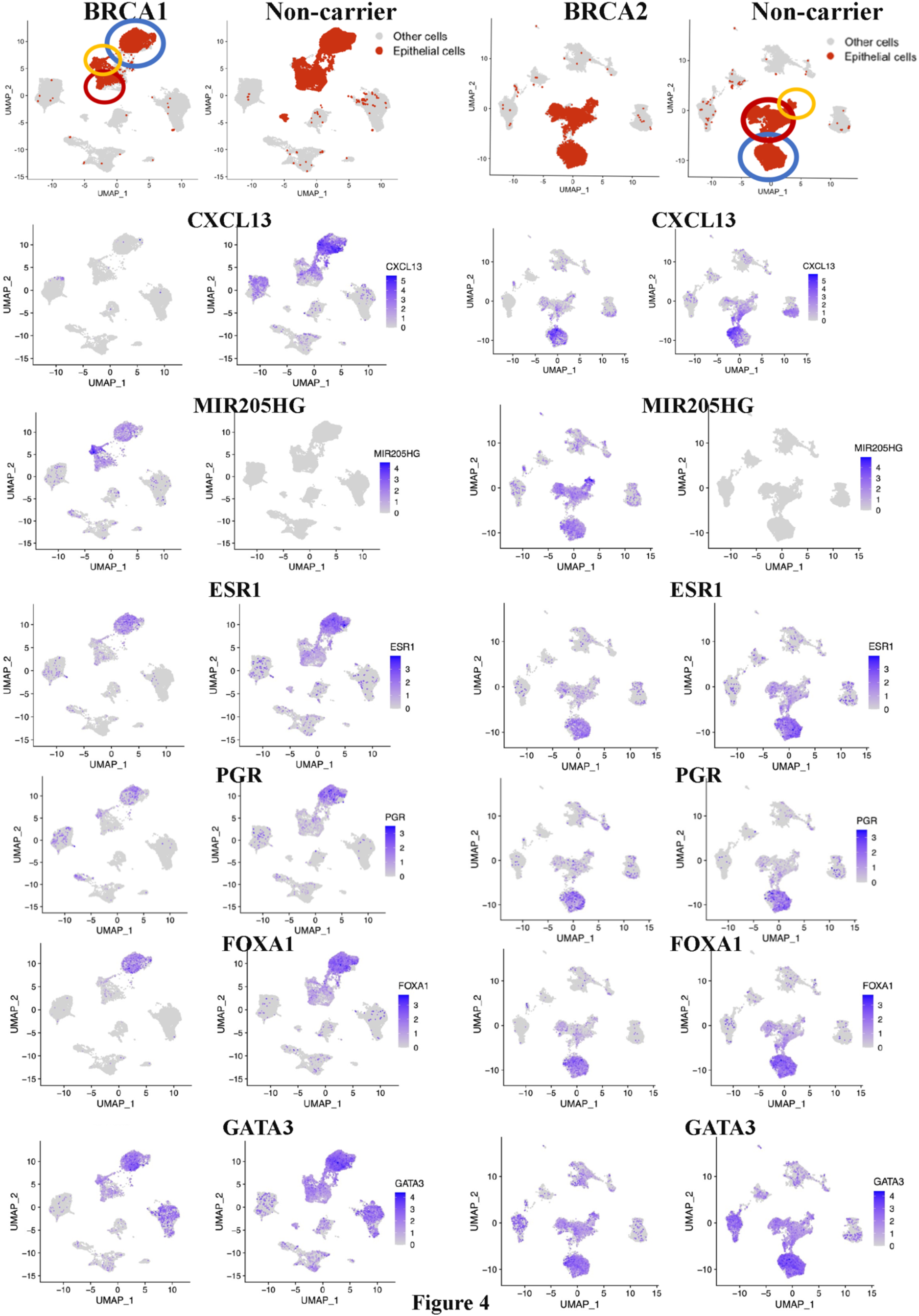
Expression patterns of select genes in breast epithelial cells of BRCA1 and BRCA2 mutation carriers compared to non-carriers. Mature luminal cells (based on ESR1, FOXA1, PGR and GATA3 expression, blue circle)), luminal progenitor/alveolar cells (based on KIT and ELF5 expression, red circle, Figure S1 for more details), and basal cell (based on TP63, ACTG2, MYL9 and OXTR expression, orange circle) populations are indicated. Note that CXCL13 expression is absent only in BRCA1 carrier derived cells. MIR205HG expression was observed only in epithelial cells of BRCA1 and BRCA2 mutation carriers.

We specifically focused our attention on levels of hormone receptors based on a previous report of antagonism of estrogen receptor (ER) function by BRCA1 (23). Levels of ESR1, which codes for ER, and its downstream target progesterone receptor (PGR), as well as androgen receptor (AR) were lower in mature luminal cells of BRCA1 mutation carriers compared to non-carriers (Figure 4 and Figure S1). However, these differences were not observed between BRCA2 mutation carriers and non-carriers. A modest decrease in the levels of transcripts corresponding to ER pioneer factors FOXA1 and GATA3 (24) were also noted between BRCA1 mutation carriers and non-carriers. However, the level of another ER co-activator, CITED1 (25), was higher in mature luminal cells of BRCA1 mutation carriers than in non-carriers (Figure S1). We also examined whether expression levels of ER responsive genes are different in BRCA1 and BRCA2 mutation carriers compared to non-carriers by focusing on three major ER target genes; PDZK1, SERPINA1, and SPDEF (19). Mature luminal cells of only BRCA1 mutation carriers expressed higher levels of these genes (Figure S2). The above results suggests that this hormonal signaling network functions differently in mature luminal cells of BRCA1 mutation but not BRCA2 mutation carriers compared to non-carriers. Other notable differences include KIT (a luminal progenitor marker) (4), ELF5 (alveolar progenitor marker) (26) and TNSF11 (also called RANKL, a target of PR) (27) (Figure S1). While a subpopulation of mature luminal cells expressed KIT in non-mutation carriers and BRCA2 mutation carriers, KIT expression was restricted to a fraction of luminal progenitors in case of BRCA1 mutation carriers. Similarly, the expression of ELF5 was more restricted to a fraction of luminal progenitors in cases of BRCA1 mutation carriers compared to others. Consistent with lower activity of PR, mature luminal cells of BRCA1 mutation carrier expressed very little TNSF11 compared to mature luminal cells of BRCA2 mutation carriers and non-carriers. Therefore, BRCA1 and BRCA2 mutations cause changes in the expression levels of specific genes.

Data presented in Figure 2 suggested that BRCA1 mutation carriers have a higher proportion of basal/stem cells compared to luminal progenitors. To further validate this observation, we determined whether expression levels of three basal cell markers are higher in epithelial cells of BRCA1 mutation carriers compared to non-carriers or BRCA2 mutation carriers. Indeed, the expression levels of ACTG2, MYL9 and OXTR, basal cell contractility genes (19), were higher in epithelial cells of BRCA1 mutation carriers compared to others (Figure S2).

### Luminal progenitors of BRCA1 and BRCA2 mutation carriers express higher levels of select genes that constitute Basal-Luminal (BL) hybrid gene signature

A recent study described a subset of alveolar cells called basal-luminal (BL) hybrid cells, which show higher levels of plasticity. It was also shown that their numbers in the breast increase with age (19). These cells carry a gene signature associated with basal-like breast cancer. Since BRCA1/2 mutation carriers display an accelerated aging phenotype (2), we compared expression of genes in BL signature in BRCA1 or BRCA2 mutated epithelial cells compared with non-carrier epithelial cells. Additionally, we verified whether genes that are shown to be differentially expressed in epithelial cells and fibroblasts of BRCA1 mutation carriers compared to non-carriers in another study are similarly differentially expressed in our dataset (21). We next examined the cell types that show differences in gene expression at individual gene level as examining gene expression differences in bulk epithelial cells of BRCA1/2 mutation carriers compared to non-carriers did not show much of a difference. BL-enriched genes are expressed in a specific subpopulation of KIT^+^ luminal progenitor cells (Figure 5). There were few differences between BRCA1 and BRCA2 as KRT6B, a breast cancer stem cell marker (28), is upregulated in luminal progenitor cells of BRCA1 mutation carriers compared to others. This is significant as KRT6B is typically expressed at higher levels in basal-like breast cancers, a type of cancer type common among BRCA1 mutation carriers (29).

**Figure 5:**
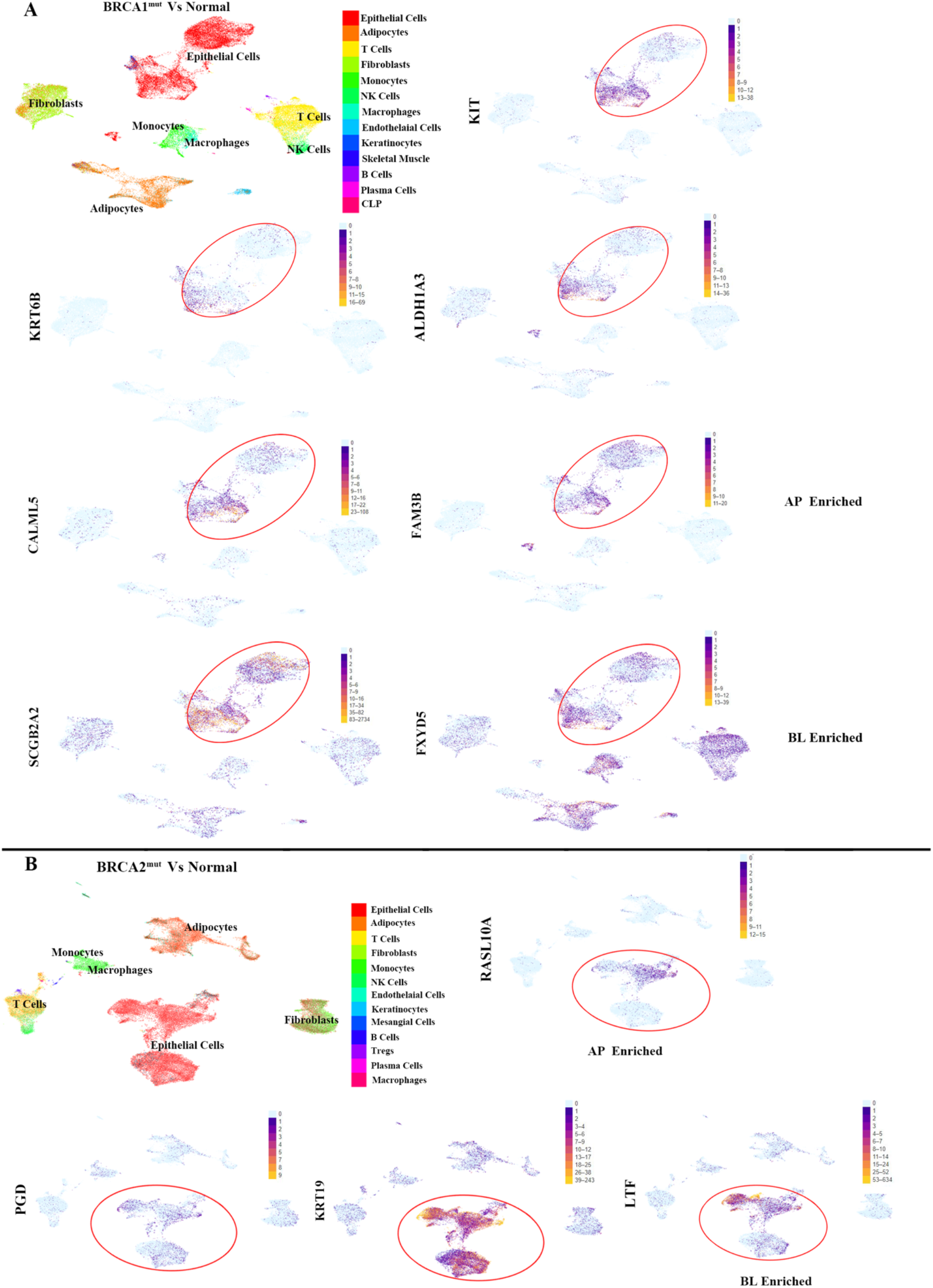
Expression levels of previously described Basal-Luminal enriched (BL) genes as well as those described to be enriched in breast epithelial cells of BRCA1 mutation carriers. A) Expression patterns of specific genes in cells from BRCA1 mutation carrier compared to non-carriers. Clusters colored light blue show no difference in expression. Alveolar progenitor (AP) cell enriched and BL cell enriched genes are indicated. KIT expression is shown to indicate epithelial cell clusters with luminal progenitor properties. Note a unique subpopulation within luminal progenitors that express higher levels of ALDH1A3 in BRCA1mutation carriers. CALML5 and FAM3B are AP enriched whereas SCGB2A2 and FXYD5 are BL enriched genes. B) Expression patterns of specific genes in cells from BRCA2 mutation carriers compared to non-carriers,

Another recent study described transcription factor networks that presumably are functionally involved in identity of mature luminal (designated as luminal-hormone responsive in the study), luminal progenitor (luminal secretory in the study), and basal cells (30). Among these transcription factors, notably elevated expression of XBP1 was observed in a subpopulation of mature luminal cells of BRCA1 and BRCA2 mutation carriers compared to non-mutation carriers (Figure S3).

### BRCA1 and BRCA2 mutations lead to activation specific signaling networks, including NF-κB, in epithelial cells

We subjected genes differentially expressed in epithelial cells of BRCA1 and BRCA2 mutation carriers compared to non-carriers to Ingenuity Pathway Analysis (IPA) to determine the effects of these mutations on basal signaling pathways. Four predominant networks are shown in Figure 6. Signaling from IKBKB, which is known to activate NF-κB (31), is elevated in both BRCA1 and BRCA2 mutation carriers. Signaling from LARP1, which links signaling from mTOR to translation of specific mRNAs (32), is also elevated in BRCA1 and BRCA2 mutation carriers. Interestingly, BRCA1 and BRCA2 mutations negatively affected signaling by cMyc, which is likely responsible for lower expression of select genes in the translational machinery. This characteristics of BRCA1 and BRCA2 mutant cells is reminiscent of embryonic diapause-like state maintained by drug tolerant cells (33). Pathways uniquely activated in BRCA1 mutated cells include BRD4, Inhibin A, HIF2A/EPAS1, and STAT3. Pathways uniquely activated in BRCA2 include CREB1, IL6, TNSF11, PDK1 and FOXO3. Upstream regulator analysis indicated specific activation of LARP1 signaling and inhibition of MYCN signaling in both BRCA1-mutant and BRCA2-mutant epithelial cells compared to non-career epithelial cells.

**Figure 6:**
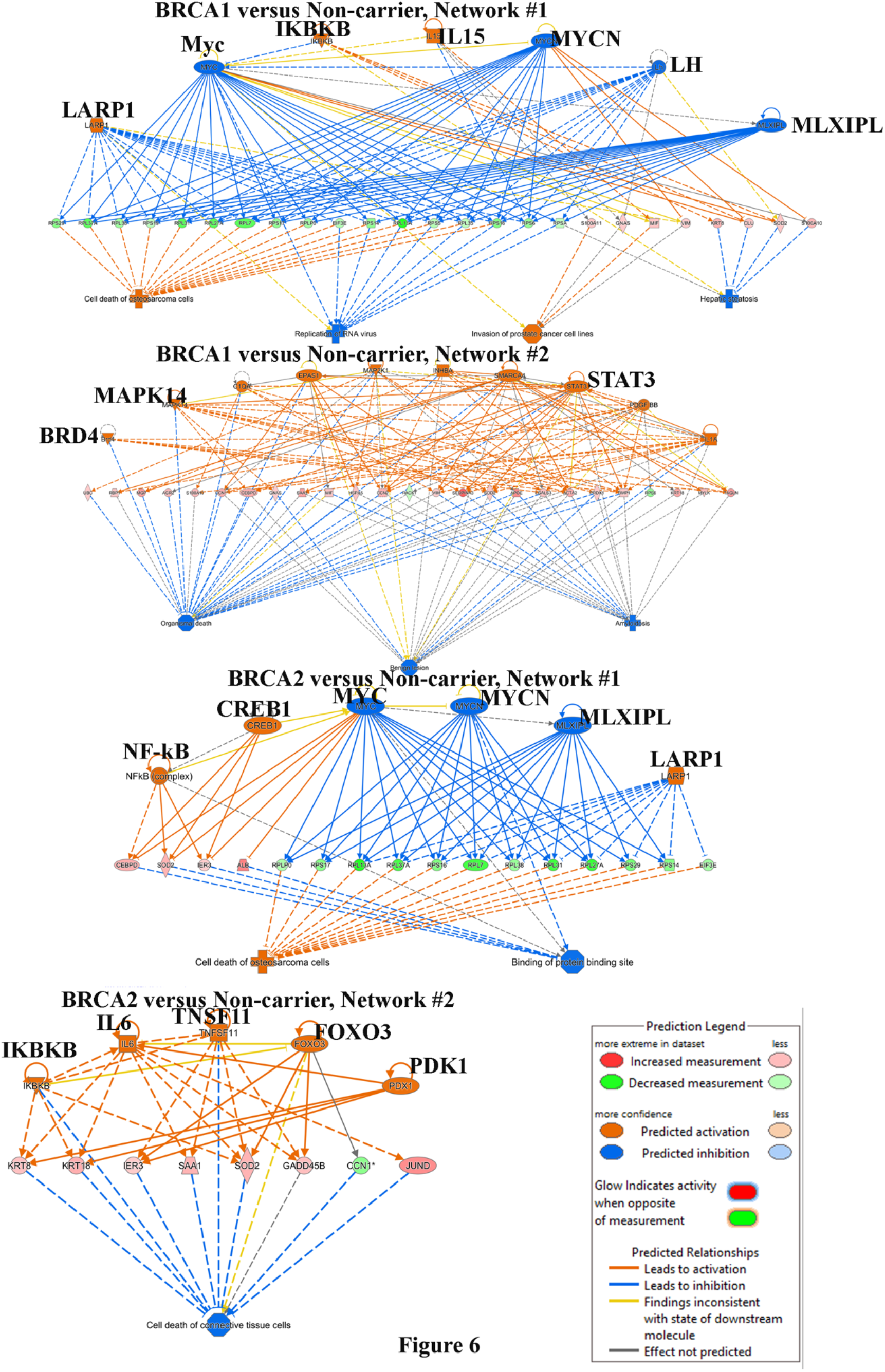
Breast epithelial cells of BRCA1 and BRCA2 mutation carriers display distinct signaling pathway activation. Signaling pathways active in epithelial cells of BRCA1 (top two) and BRCA2 (bottom 2) compared to epithelial cells of non-carriers are shown.

### Distinct differences in immune cell composition of breast tissues of BRCA1 and BRCA2 mutation carriers compared to non-carriers

Since ∼20% of cells sequenced were immune cells in each group (Table S2), we examined whether there are any qualitative differences in immune cell types that infiltrate breast tissues between the three groups. Overlay analysis of T cells and macrophages from BRCA1 or BRCA2 carriers over non-carriers showed specific differences (Figure S4). For example, breast tissues from BRCA1 and BRCA2 mutation carriers are enriched for Interleukin 17 receptor-positive (IL17R^+^), Granulysin-positive (GNLY^+^) effector T cells, and Granzyme K-positive (GZMK^+^) cytotoxic T cells, which are enriched in the microenvironment of Triple Negative Breast Cancer (TNBCs) (34–36), and pro-inflammatory Triggering Receptor Expressed in Myeloid Cells 2-positive (TREM2^+^) macrophages, which are enriched in the microenvironment of multiple tumor types with pro-tumorigenic activities (37–39). We also observed elevated levels of C-X-C Chemokine Receptor 4-positive (CXCR4^+^) T cells, monocytes/macrophages and a subpopulation of luminal progenitor cells in BRCA1 mutation carriers. CXCR4 positivity was modest in case of BRCA2 mutation carriers. Similarly, Interferon gamma (IFNG) expressing T cells are enriched in BRCA1/2 mutation carriers. These results are consistent with data described in the recent single cell study of reduction mammoplasty samples (30). Thus, it is possible that the immune microenvironment in the breast tissues of BRCA1 and BRCA2 carriers is inherently enriched for pro-tumorigenic immune cells.

### Immortalized breast epithelial cell lines from BRCA1/2 mutation carriers show elevated NF-κB activation

To determine whether pathways activated in BRCA1 and BRCA2 mutant epithelial cells compared to epithelial cells from non-carriers, as suggested by scRNA-seq, are carried over to immortalized cells and are observed in vitro, we established immortalized cell lines from

BRCA1/2 mutation carriers. The immortalized cell line from a BRCA1 mutation carrier has been described previously (12) and we created a new cell line from a BRCA2 mutation carrier. We also overexpressed PIK3CA^H1047R^ mutant in these cell lines to compare the effects of oncogene overexpression on cell phenotype and signaling networks. We selected PIK3CA^H1047R^ mutant as an oncogene because mutation in the *PIK3CA* gene is the second most common mutation in breast cancer after *TP53* (40). Flow cytometry characterization of these cells demonstrated that immortalized cells from BRCA1 mutation carrier are enriched for both basal and luminal progenitors, whereas cells from BRCA2 mutation carrier are predominantly luminal progenitors (Figure 7A). Interestingly, PIK3A^H1047R^ overexpression resulted in a significant increase in EpCAM expression in the cell line derived from the BRCA1 mutation carrier (CD49f^+^/EpCAM^+^ cells increased from 12% to 36% upon PIK3CA^H1047R^ overexpression). This is intriguing in light of recent observation that EpCAM^high^ cells but not cells that have undergone stable epithelial to mesenchymal transition are the highly metastatic subpopulation of cancer cells (41). PIK3CA^H1047R^ mutant also increased the number of CD44^+^/CD24^+^, CD271^+^/EpCAM^+^ and CD44^+^/EpCAM+ cells at the expense of CD44^+^/CD24^-^, CD271^+^/EpCAM^-^ and CD44^+^/EpCAM^-^ cells, respectively, in case of BRCA1 mutation carrier. Thus, BRCA mutation status influences the ability of mutant PIK3CA to alter the differentiation pathway of breast epithelial cells and potentially influence metastatic properties of cancer cells.

**Figure 7:**
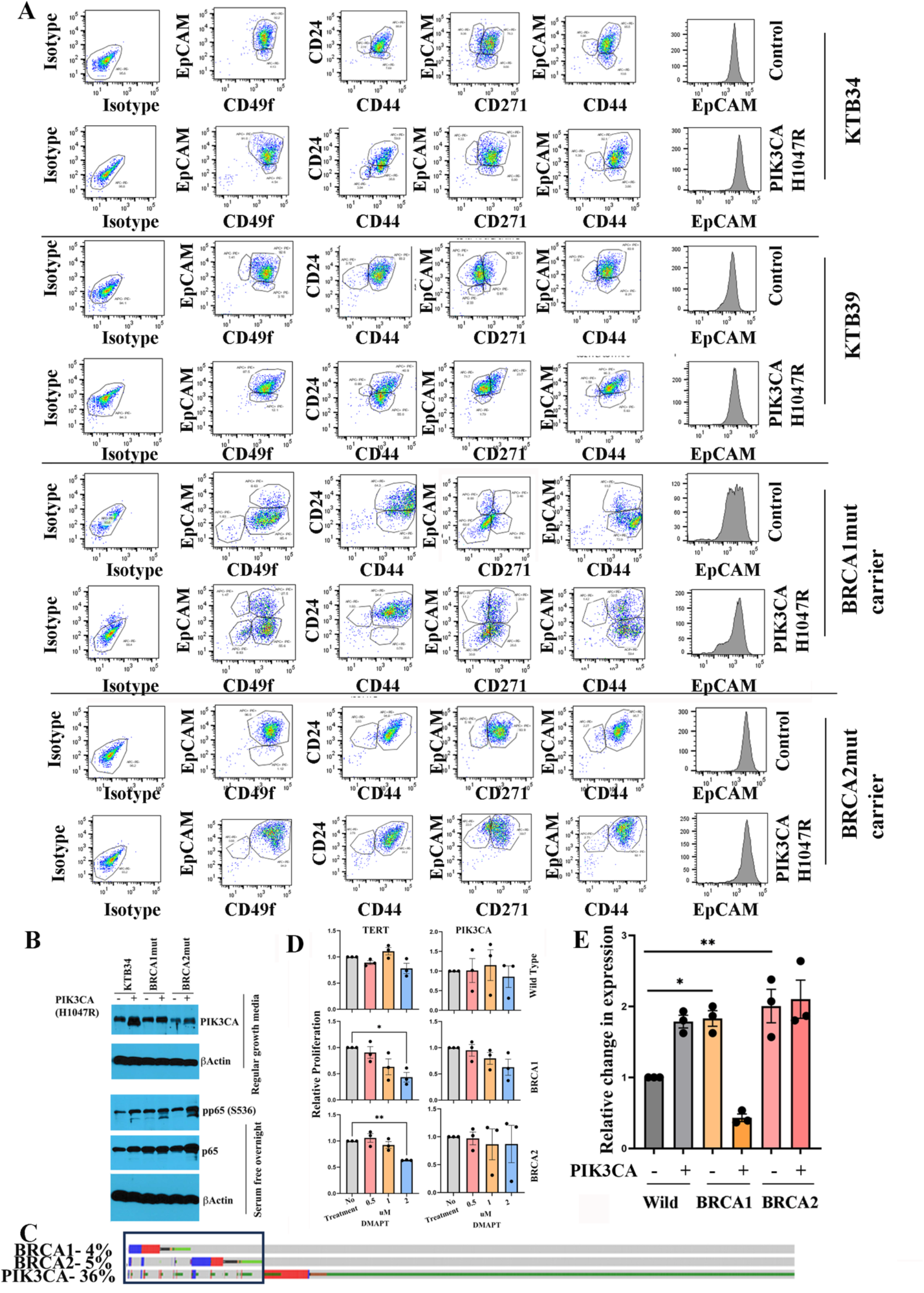
Immortalized BRCA1 or 2 mutant cells display elevated NF-κB activity. A) PIK3CA^H1047R^ distinctly influences differentiation properties of immortalized BRCA1 and BRCA2 mutation carriers. Vector control or PIK3CA^H1047R^ expressing breast epithelial cell lines from non-carriers (KTB34 and KTB39) and BRCA1 or BRCA2 mutation carriers were stained with indicated antibodies and characterized by flow cytometry (n=3). Representative data are shown. PIK3CA^H1047R^ robustly increased EpCAM expression and increased differentiated phenotype of BRCA1 mutant cells. CD49f^+^/EpCAM+ cells increased from 12% to 36% whereas CD49f^+^/EpCAM^-^ cells decreased from 80% to 34% upon PIK3CA^H1047R^ overexpression. B) BRCA1 mutant cells display elevated phosphorylation of p65, a NF-κB subunit, which indirectly suggests activation of NF-κB. Expression levels of PIK3CA in cell transduced with PIK3CA mutant virus are shown. Regular growth media condition had to be used to detect PIK3CA mutant overexpression because robust induction of endogenous PIK3CA upon serum starvation. An antibody that preferentially recognizes PIK3CA^H1047R^ mutant was used in the western blot. C) Approximately 30% of breast cancers in BRCA1 or BRCA2 mutation carriers carry PIK3CA mutations. Data were generated using cBioportal. D). Immortalized BRCA1 or BRCA2 mutant cells but not cells from non-carriers are sensitive to DMAPT. PIK3CA overexpression reduced sensitivity to DMAPT. E) Immortalized cell lines from BRCA1 or BRCA2 mutation carriers express higher levels of MIR205HG. Note that PIK3CA^H1047R^, which robustly induced differentiation of BRCA1 mutant cells, reduced MIR205HG levels in these cells. P values, *<0.05; **<0.01.

scRNA-seq predicted constitutive activation of NF-κB in BRCA1 and BRCA2 mutant cells. To validate this observation, we measured the phosphorylation status (S536) of the p65 subunit of NF-κB. S536 phosphorylation of p65 by kinases such as IκBβ leads to its increased activity and this phosphorylation status is an indirect measure of its activity (42). Basal phospho-p65(S536) levels were higher in immortalized BRCA1 mutant and BRCA2 mutant cells compared to cells from non-carrier (Figure 7B). These observations may be clinically relevant as ∼30% of breast cancers in BRCA1 or BRCA2 mutation carriers carry *PIK3CA* mutations (Figure 7C). To determine whether BRCA1 or BRCA2 mutant cells show dependency on NF-κB for survival, we measured the sensitivity of these cells to dimethylaminoparthenolide (DMAPT), a NF-κB inhibitor we described previously (43). Indeed, immortalized BRCA1 and BRCA2 mutant cells but not cells from non-carriers were sensitive to DMAPT (Figure 7D). Interestingly, PIK3CA^H1047R^ overexpression reversed the dependency on NF-κB. These results indicate that germline mutations coupled with the type of oncogenic mutation determine sensitivity of cells to specific therapies.

scRNA-seq data suggested elevated MIR205HG expression in BRCA1 and BRCA2 mutant breast epithelial cells compared to cells from non-carrier cells. Because of the emerging role of MIR205HG in cellular processes such as prostate basal cell differentiation (44), its regulation by super-enhancer (45), and microRNA generated from its transcripts targeting BRCA1 (46), we verified its expression in immortalized cells from non-carriers and BRCA1 and BRCA2 mutation carriers. Indeed, MIR205HG levels are ∼2-fold higher in BRCA1 and BRCA2 mutant cells compared to cells from non-carriers (Figure 7E). Consistent with effects of PIK3CA^H1047R^ on differentiation of BRCA1 mutant cells, its overexpression reduced MIR205HG in only BRCA1 mutant cells. Thus, MIR205HG could be one of the previously uncharacterized downstream mediators of the effects of BRCA1 and BRCA2 mutation.

### BRCA1 and BRCA2 mutant cells overexpressing PIK3CA^H1047R^ are tumorigenic in NSG mice

We recently reported that overexpression of PIK3CA^H1047R^ is insufficient to induce transformation of immortalized luminal breast epithelial cells but overexpression of PIK3CA^H1047R^ along with SV40-T/t antigens generate transformed cells that develop non-metastatic adenocarcinomas in NSG mice (13). SV40-T/t antigens inactivate multiple tumor suppressor pathways including p53, retinoblastoma, PP2A; and deregulate multiple DNA damage signaling and repair pathways (47,48). Because BRCA1 and BRCA2 mutations also lead to impaired DNA damage response, we next examined whether BRCA1 and BRCA2 mutations can substitute for SV40-T/t antigens in PIK3CA^H1047R^-mediated tumorigenesis. Towards this goal, we injected immortalized BRCA1 or BRCA2 mutant cells expressing PIK3CA^H1047R^ into the mammary fat pad of NSG mice. Five animals per group were injected and the experiment was done twice. Tumor growth patterns are described in Figure 8A. Animals were sacrificed ∼3 months post-injection. In both series of experiments, no tumors developed when cells overexpressing HRAS^G12V^ or SV40-T/t antigens were used but the combination of both was effective in generating tumors. In the first series, three out of five animals injected with BRCA1 or BRCA2 mutant cells overexpressing PIK3CA^H1047R^ developed tumors. Four out five and all animals injected with BRCA1+PIK3CA^H1047R^ and BRCA2+PIK3CA^H1047R^ cells developed tumors in the second round of experiments. H&E staining patterns, histologic characterization and expression of luminal markers ERα and GATA3, and keratins CK14 and CK19 of two tumors in each category are shown in Figure 8B. Although BRCA1 mutation carriers rarely develop ER^+^ tumors, in our model, tumors were heterogenous with a fraction of tumor cells expressing ERα and GATA3. As confirmed by pathologist, all tumors were invasive ductal carcinomas. With respect to BRCA2, most tumors were cystic and basal cell carcinomas with unique keratin expression pattern. However, these tumors still expressed the luminal marker GATA3. We did not observe any lung metastasis. Overall, these results suggest that BRCA1 and BRCA2 mutations can effectively substitute the need for SV40-T/t antigens to achieve transformation of breast epithelial cells by an oncogene.

**Figure 8:**
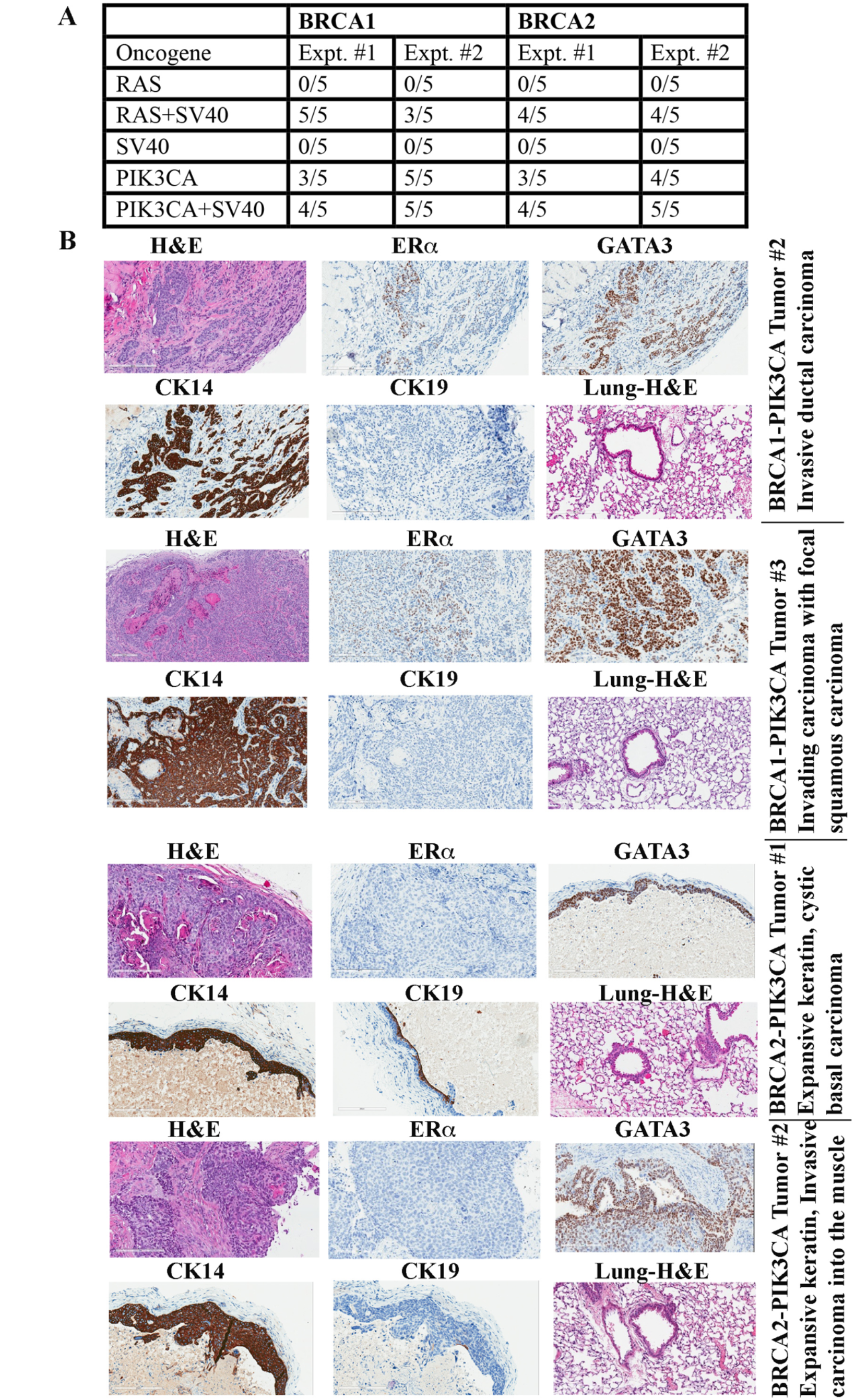
Immortalized breast epithelial cells from BRCA1 or BRCA2 mutation carriers overexpressing PIK3CA^H1047R^ mutant generate tumors in NSG mice. A) Frequency of tumor formation. Number of animals injected and those that developed tumors in each of the experiments are shown. For unknown reasons, tumors in BRCA1+PIK3CA^H1047R^ cells injected animals in the second experimental series were flatter and appeared only after 12 weeks of tumor cell injection but were histologically breast epithelial cell-derived tumors. B) Immunohistochemical characterization of tumors. Tumors developed from BRCA1 mutant cells show heterogenous expression patterns of luminal markers ERα and GATA3. Tumors developed from BRCA2 mutation carriers are ERα-negative but expressed variable levels of GATA3.

## Discussion

Although susceptibility to breast cancer in BRCA1 and BRCA2 mutation carriers has been known for several decades, mechanisms responsible for this susceptibility are just beginning to be identified, largely due to recent advances in single cell technologies. Three recent studies have used single cell RNA/Protein technologies to identify differences in breast cell types between BRCA1/2 mutation carriers and non-carriers. Gray et al suggested that BRCA2 mutation carriers contain a lower number of progesterone receptor (PR)-positive hormone responsive cells compared to non-carriers; and their study was done at single cell protein level using CyTOF (19). In this study, we did not observe significant differences in levels of PR+ cells between BRCA2 mutation carriers compared to non-carriers based on mRNA expression (Figure 4). However, we observed fewer PR+ cells in BRCA1 mutation carriers than non-carriers. The second study by Nee et al found differences in BL intermediate progenitor cell types and stromal cells in BRCA1 mutation carriers compared to non-carriers (21). Consistent with those results, we found elevated expression of several genes associated with BL progenitor phenotype in BRCA1 mutation carriers compared to non-carriers (Figure 5). Differences between BRCA2 mutation carriers and non-carriers are less evident. The third study found elevated basal progenitor cells in BRCA1 mutation carriers compared to non-carriers although number of non-carriers in the study was only three samples (7). That study reported high levels of ALDH1A3^+^ cells in BRCA1 mutation carriers, similar to the second report (21). We also observed elevated ALDH1A3 expression in BRCA1 but not in BRCA2 mutation carriers compared to non-carriers (Figure 5). Thus, elevated expression of ALDH1A3 in BRCA1 mutation carriers is consistent across multiple studies. Since ALDH1A3 is a normal stem and cancer stem cell marker (49), these results suggest that breast epithelial cells in the normal breasts of BRCA1 mutation careers have inherently higher stem cell activity. Collectively, all four studies including ours suggest an effect of BRCA1 mutations on basal-luminal hybrid phenotype and acquisition of ALDH1A3 positivity.

Previous studies in mouse models have shown the effects of BRCA1 mutation on NF-κB activation (15). Gray et al also suggested that increased NF-κB activity could be responsible for basal-luminal hybrid cell plasticity of BRCA1-mutant epithelial cells (19). Nee et al observed elevated levels of IκBα, an inhibitor of NF-κB, in normal breast epithelial cells compared to BRCA1 mutant breast epithelial cells, indirectly suggesting increased NF-κB activity in BRCA1 mutant cells (21). Our studies also clearly show elevated NF-κB signaling in both BRCA1 and BRCA2 mutant cells (Figure 7). How NF-κB remains active in BRCA1 and BRCA2 mutant cells is unknown. A recent study reported transcriptional reprogramming in BRCA1-deficient ovarian cancer cells, which leads to cell-intrinsic inflammation through activation of stimulator of IFN genes (STING) (50). Increased STING activity leads to chronic inflammation through the NF-κB pathway (51,52). It is, therefore, possible that even “normal” cells in BRCA1 and BRCA2 mutation carriers have elevated basal STING activity and chronic inflammatory phenotype. As STING agonists are currently being tested in preclinical models to improve immunotherapy (53), STING antagonists may need to be developed as chemoprevention agents for BRCA1/2 mutation carriers.

BRCA1 and BRCA2 are involved in different steps of the same homologous recombination-mediated DNA repair pathway (54). However, breast tumors in BRCA1 and BRCA2 mutation carriers generally show different histopathology. While BRCA1 mutation carriers typically develop basal-like breast cancers, breast cancers in BRCA2 mutation carriers are much more heterogenous (55). While the risk of contralateral breast cancer in BRCA1 mutation careers decreases after menopause, incidence increases in BRCA2 mutation carriers. Despite different pathophysiology, we did not observe distinct differences in epithelial cell populations of BRCA1 and BRCA2 mutation carriers. We found only 26 genes being differentially expressed between epithelial cells of BRCA1 mutation carriers compared to BRCA2 mutation carriers (Table S5). There was an even lower number of differences in stromal fibroblasts between BRCA1 and BRCA2 mutation carriers (12 genes). One major difference we found was the degree to which BRCA1 epithelial cells express genes associated with basal-luminal hybrid phenotype and plasticity of epithelial cells. It is possible that enhanced plasticity of BRCA1 mutant epithelial cells make these cells more susceptible to basal-like breast cancers, whereas limited plasticity makes tumors in BRCA2 mutation carriers similar to sporadic breast cancers. Although individual gene level differences between BRCA1 and BRCA2 epithelial cells were minor, we did observe several differences in signaling pathways (Figure 6). For example, PDK1 is uniquely activated in BRCA2 mutation carriers. Specific activation of PDK1 in BRCA2 mutation carriers is interesting, as this kinase has recently been shown to confer resistance to CDK4/6 inhibitors in ER^+^ breast cancer cell lines, and inhibitors targeting this kinase are under development (56). Models created here should help to further evaluate this possibility.

Recent studies have shown that a number of cancer driver mutations are found in normal tissues suggesting that these driver mutations alone are insufficient to initiate cancer (57). Mutations in genes such as *ARID1A, PIK3CA, ERBB2, FAT1, KMT2D*, and *TP53* are found in several cancer-free organs including bladder, colon, liver, and endometrium. Similarly, *PIK3CA^H1047R^* mutation is found in 22% of benign breast biopsies that did not progress to cancer within a year of tissue collection and in 19% of cases which did progress to cancer (58). These observations suggest that *PIK3CA* mutation alone is not sufficient to initiate breast cancers and mutations that co-occur with it are needed to initiate breast cancer. We and others have shown that efficient transformation of primary breast epithelial cells requires a combination of three oncogenes: hTERT, SV40-T/t antigens and mutated H-RAS or PIK3CA (13,59). The observation in this study that BRCA1 or BRCA2 mutations can substitute for SV40-T/t antigens for transformation by hTERT+PIK3CA^H1047R^ suggests that aberration in signaling molecules that co-operate with BRCA1 or BRCA2 in DNA repair pathways could be the second mutation along with a *PIK3CA* mutation needed to initiate breast cancer. Future studies focused on identifying such mutations would pave the way to identify minimum oncogenic mutations that lead to breast cancer initiation.

## Supporting information

Uncropped Western blot images

## Acknowledgements

We thank IUSCCC tissue bank providing all tissues utilized in the study and Flow cytometry core for the analysis. These cores are supported by the Cancer Center Support grant P30CA082709. We also thank Immunohistochemistry core for analysis of tumors.

## Author contributions

Conception and design: HN

Development of methodology: PBN, AK, DC, KB, HM, CH

Acquisition of data: PBN, AK, DC, KB, HM, CH, PCM, XX, GS, HN

Analysis and interpretation of data: PBN, AK, DC, HG, YL, GS, AMS, HN

Administrative, technical, or material support: AMS, HN

Writing and review of the manuscript: PBN, AK, AMS and HN

Study supervision: HN

## Data and Material availability

All data needed to evaluate the conclusions in the paper are present in the paper and/or in the supplementary materials. Sequence data have been submitted to publicly available databases and accession numbers are indicated in the Materials and Methods. Requests for reagents including cell lines should be submitted to HN.

