## Supplementary Tables 1 and 2 and S1-S4 for "BRCA1 and BRCA2 germline mutations driven signaling pathway alterations are sufficient to initiate breast tumorigenesis by the PIK3CA^H1047R^ oncogene"

**Supplementary information**

**Table S1:** Information on samples used for single cell studies. All tissues are taken from the at the time of surgery as a part of treatment. Samples 1-4 are BRCA1 and 5-8 are BRCA2. In case of sample 4, tissues from left and right breasts were sequenced separately.

| Case number | Age | Race | Source | Histology | Diagnosis/Treatment |
|---|---|---|---|---|---|
| 1 | 33 | White | Normal-adjacent | Normal | Diagnosed with inflammatory breast cancer |
| 2 | 37 | White | Normal-adjacent | Normal | History of Triple negative breast cancer |
| 3 | 33 | White | Contralateral | Normal | Fibrocystic and ductal hyperplasia, preoperative chemo |
| 4 | 45 | Black | Normal adjacent and Normal | Normal | Stage I TNBC (T1cN0), partial mastectomy followed by chemo. Prophylactic surgery three months after chemo. |
| 5 | 42 | White | Normal-adjacent | Normal | ER-/PR+ tumor |
| 6 | 67 | White | Normal | Normal | ER+ tumor 20 years before bilateral mastectomy |
| 7 | 42 | Black | Normal-adjacent | Normal | Metaplastic squamous cell carcinoma-TNBC. Prior chemotherapy |
| 8 | 39 | White | Normal-prophylactic | Normal | Fibrocystic changes and fibro adenomas |

**Table S2: Number and percentage of different cell types in different tissue types used in this study.** ND= not detected

| Cell type | Non-carrier | BRCA1 | BRCA2 |
|---|---|---|---|
| **Epithelial cells** | 15577 (42.3%) | 3670 (21.3%) | 10498 (41.9%) |
| **Endothelial cells** | 446 (1.2%) | 605 (3.5%) | 214 (0.9%) |
| **Fibroblasts** | 4559 (12.4%) | 2537 (14.7%) | 2074 (8.3%) |
| **Adipocytes** | 5986 (15.3%) | 4397 (25.5%) | 9067 (36.2%) |
| **T cells** | 6217 (16.9%) | 3788 (22%) | 2023 (8.1%) |
| **Monocytes** | 2024 (5.5%) | 568 (3.3%) | 623 (2.5%) |
| **Macrophages** | 860 (2.3%) | 199 (1.2%) | 0 (0%) |
| **NK cells** | 891 (2.4%) | 784 (4.6%) | 277 (1.1%) |
| **Keratinocytes** | 34 (0.1%) | 386 (2.2%) | 172 (0.7%) |
| **Plasma cells** | 69 (0.2%) | 40 (0.2%) | 17 (0.1%) |
| **B cells** | 115 (0.3%) | 31 (0.2%) | 8 (0.03%) |
| **CLP** | 24 (0.1%) | 18 (0.1%) | ND |

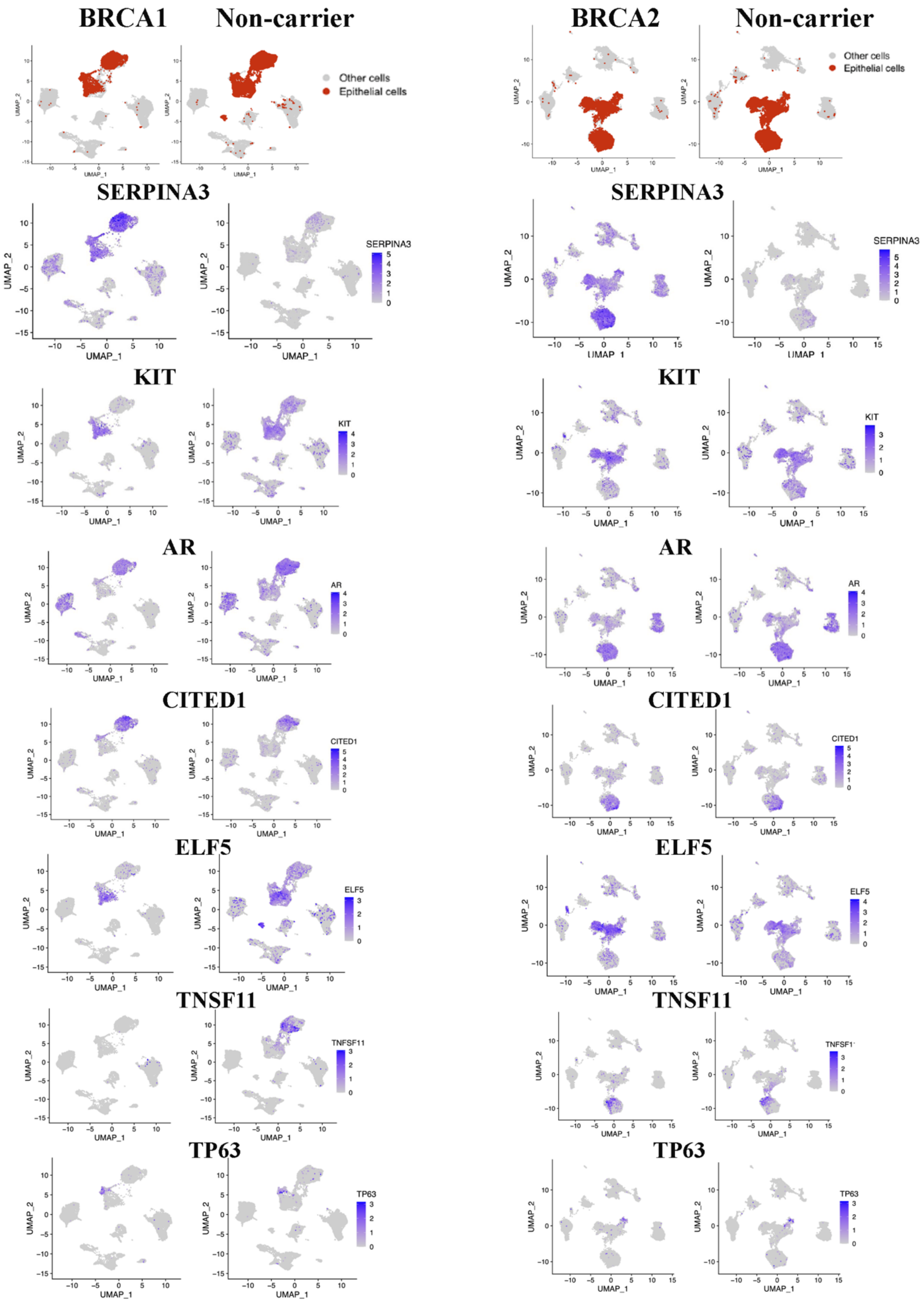
BRCA1
Non-carrier
BRCA2
Non-carrier
Other cells
Epithelial cells
SERPINA3
KIT
AR
CITED1
ELF5
TNSF11
TNFSF11
TP63
UMAP_1
UMAP_2

**Figure S1**

**Figure S1: Expression patterns of several epithelial cells enriched/specific genes in BRCA1 or BRCA2 mutation carriers compared to non-carrier.** KIT, ELF5 and TP63 expressing cells are considered to present luminal progenitor, alveolar and basal cells, respectively.

**A** **BRCA1$^{mut}$ Vs Normal**

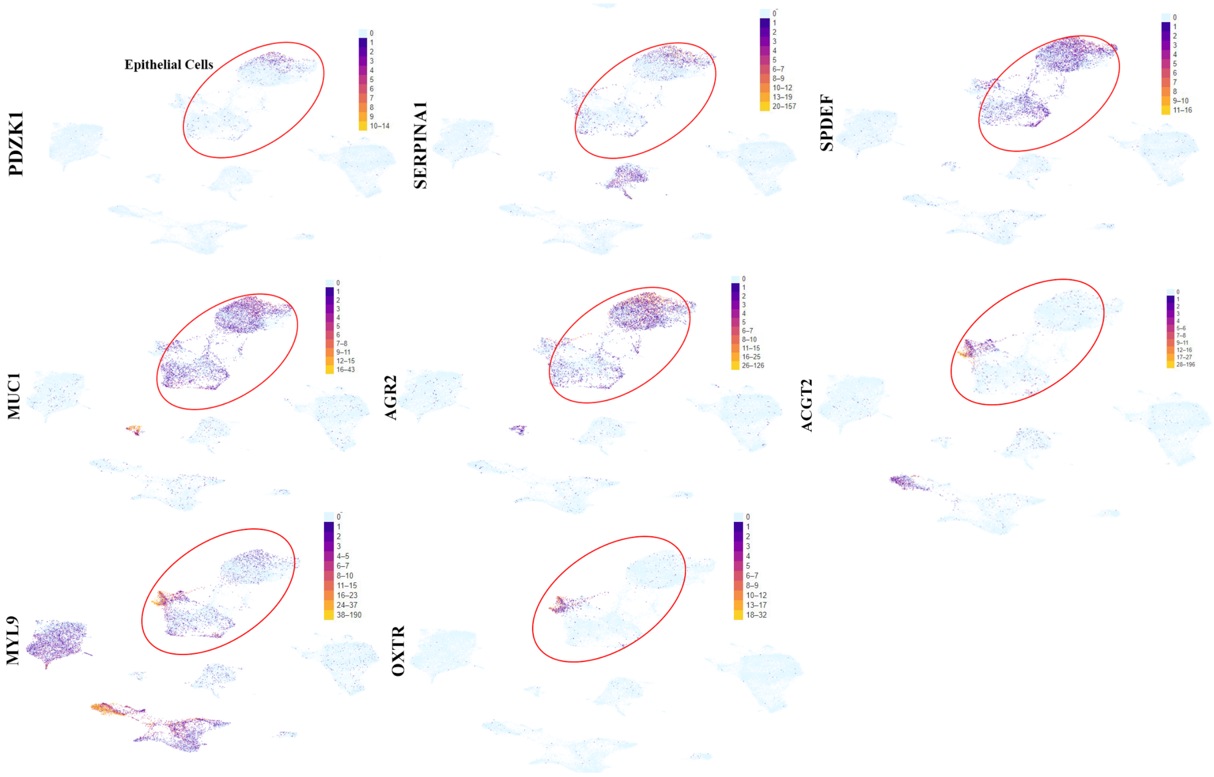


**BRCA2$^{mut}$ Vs Normal**

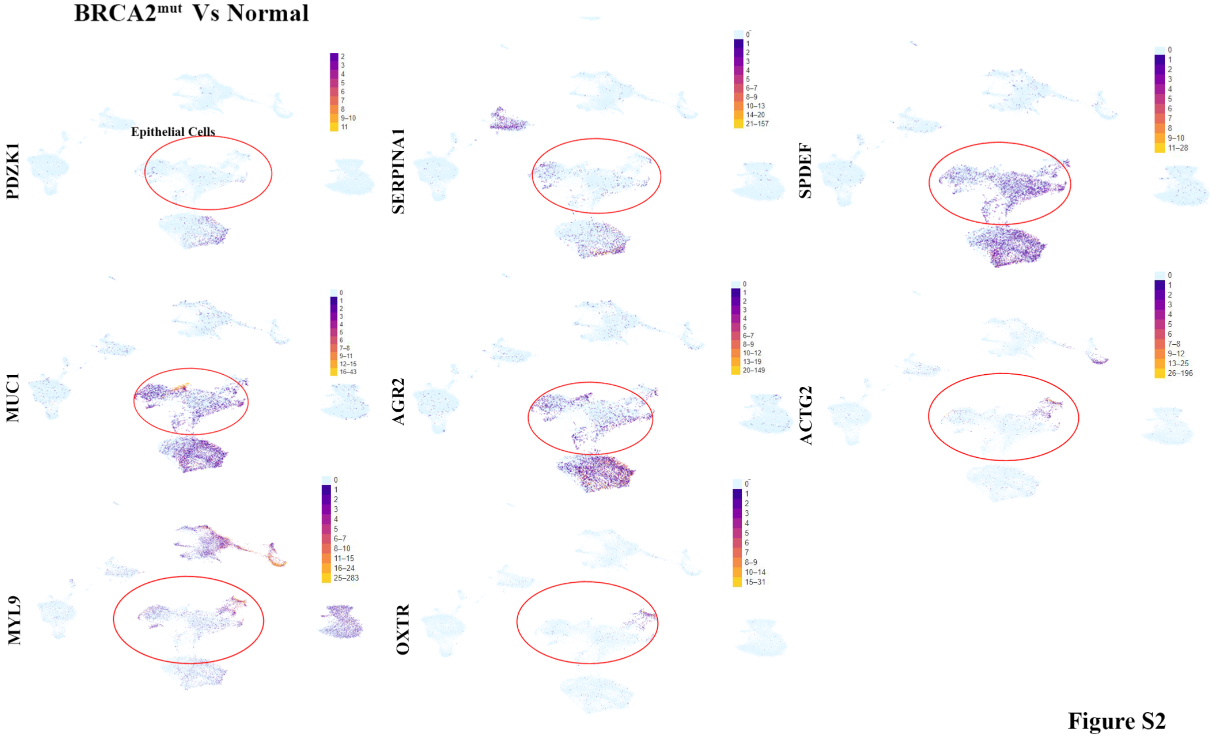


**Figure S2**

**Figure S2: Expression differences of select ERα-responsive and hormone sensitive cell marker genes in BRCA1 or BRCA2 mutation carriers compared to non-carriers.**

SERPNA1, PDZK1 and SPDEF are ERα target genes, whereas ARG2 and MUC1 and SERPNA1 are markers of hormone responsive genes.

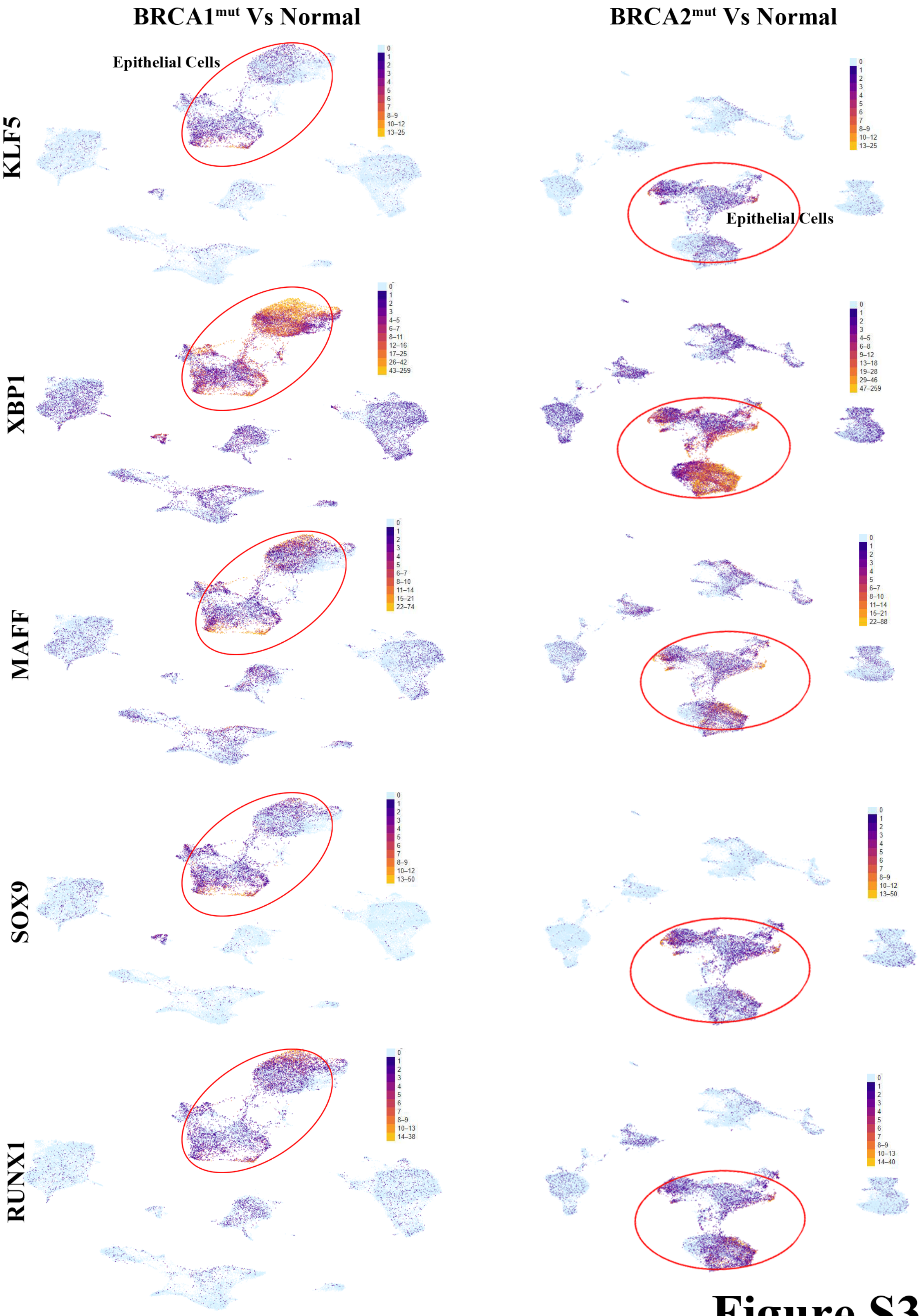
BRCA1mut Vs Normal
BRCA2mut Vs Normal
Epithelial Cells
Epithelial Cells
KLF5
XBP1
MAFF
SOX9
RUNX1

### Figure S3

**Figure S3: Expression levels of top transcription regulators in major epithelial subclusters (mature luminal, luminal progenitors and basal/stem cells).** Transcription regulators described by Kumar et al in their Extended Figure 1 [1] were evaluated in our data set and only those that showed difference is expression are shown. KLF5 (luminal progenitor/luminal secretory), XBP1 (mature luminal/luminal hormonal), MAFF (basal), SOX9, (luminal progenitor/luminal secretory), RUNX1 (mature luminal/luminal hormonal) showed variability in expression between BRCA1/2 mutant carriers compared to non-carriers with highest difference in XBP1 expression.

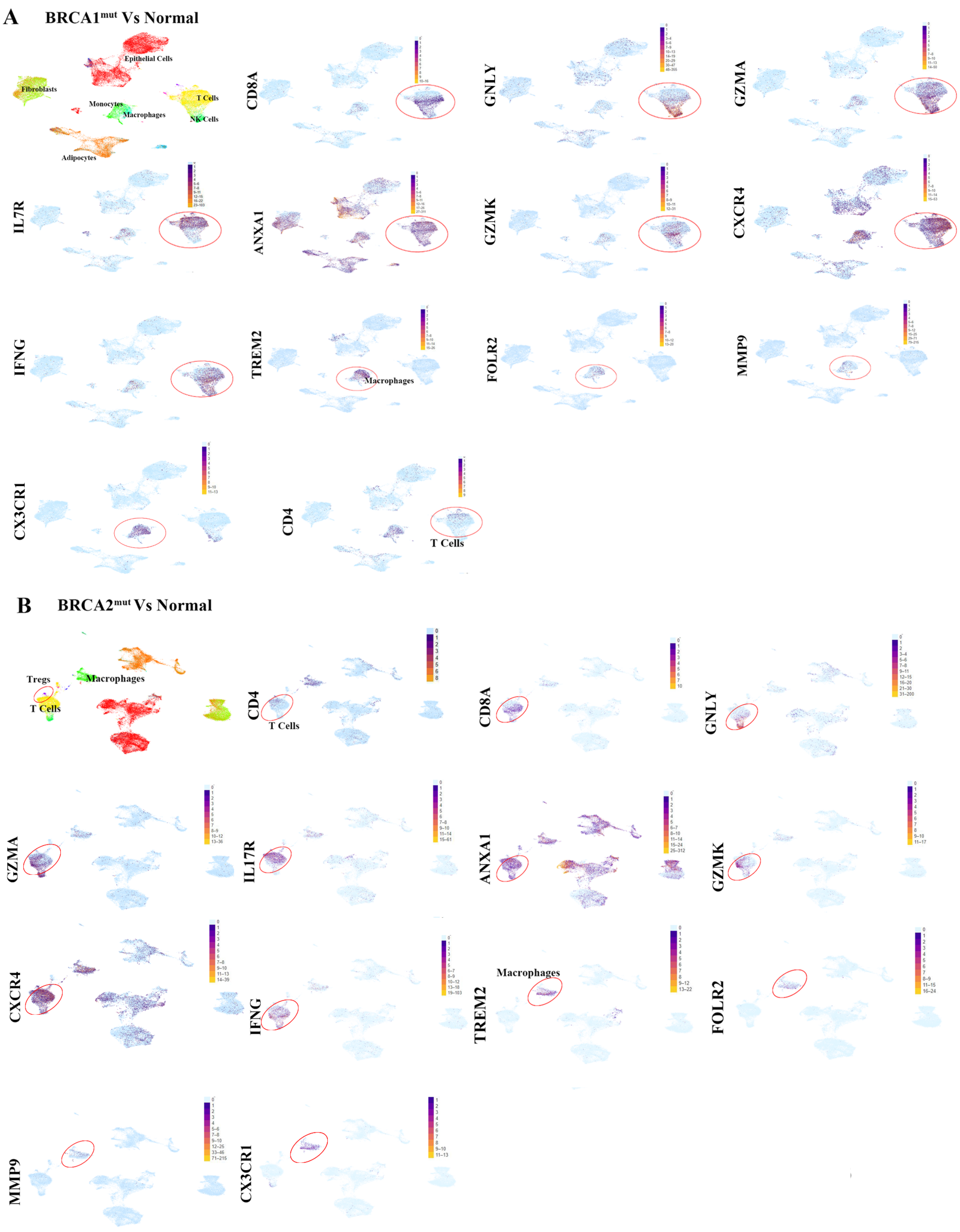

A
BRCA1mut Vs Normal
Epithelial Cells
Fibroblasts
T Cells
Monocytes
Macrophages
NK Cells
Adipocytes
CD8A
GNLY
GZMA
IL7R
ANXA1
GZMK
CXCR4
IFNG
TREM2
Macrophages
FOLR2
MMP9
CX3CR1
CD4
T Cells
B
BRCA2mut Vs Normal
Tregs
Macrophages
T Cells
CD4
T Cells
CD8A
GNLY
GZMA
IL17R
ANXA1
GZMK
CXCR4
IFNG
TREM2
Macrophages
FOLR2
MMP9
CX3CR1


**Figure S4**

**Figure S4: Expression pattern of T cells and macrophage associated genes in BRCA1 or BRCA2 mutation carriers compared to non-carriers.** Cell cluster identity depicted in Figure 5 of the main manuscript is reproduced here to allow proper assessment of results.

**Table S3: Gene expression differences in epithelial cells, endothelial cells and fibroblasts of BRCA1 mutation carrier compared to non-carrier.**

**Table S4: Gene expression differences in epithelial cells, endothelial cells and fibroblasts of BRCA2 mutation carrier compared to non-carrier.**

**Table S5: Gene expression differences in epithelial cells, endothelial cells and fibroblasts of BRCA1 mutation carrier compared to BRCA2 mutation carrier.**


1. Kumar T, Nee K, Wei R*, et al.* A spatially resolved single-cell genomic atlas of the adult human breast. Nature 2023; 10.1038/s41586-023-06252-9.
