## Supplementary material for "BRCA1 and BRCA2 germline mutations driven signaling pathway alterations are sufficient to initiate breast tumorigenesis by the PIK3CA^H1047R^ oncogene": Uncropped Western blot images

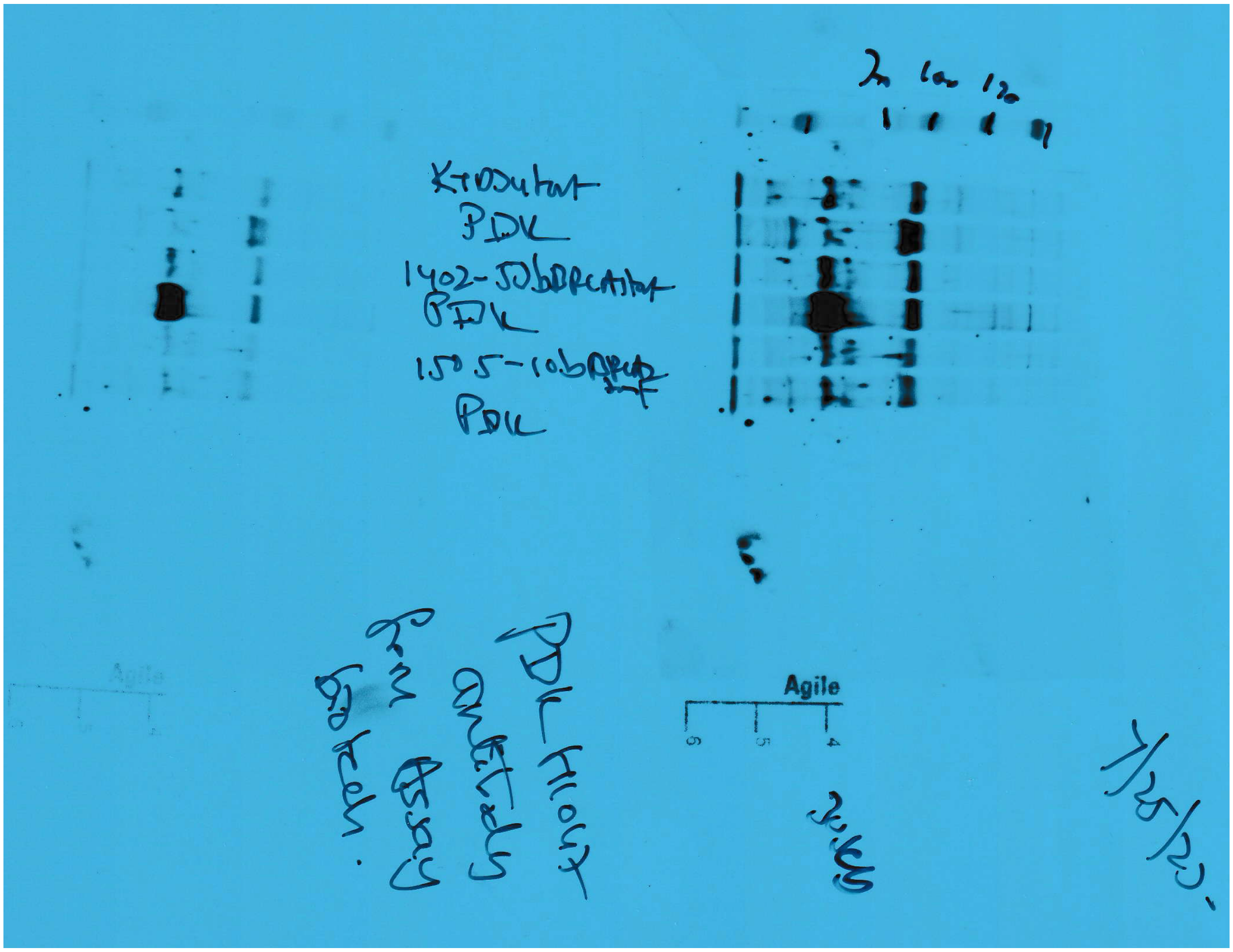
PDK
PDK
PDK
Agile
7/25/20

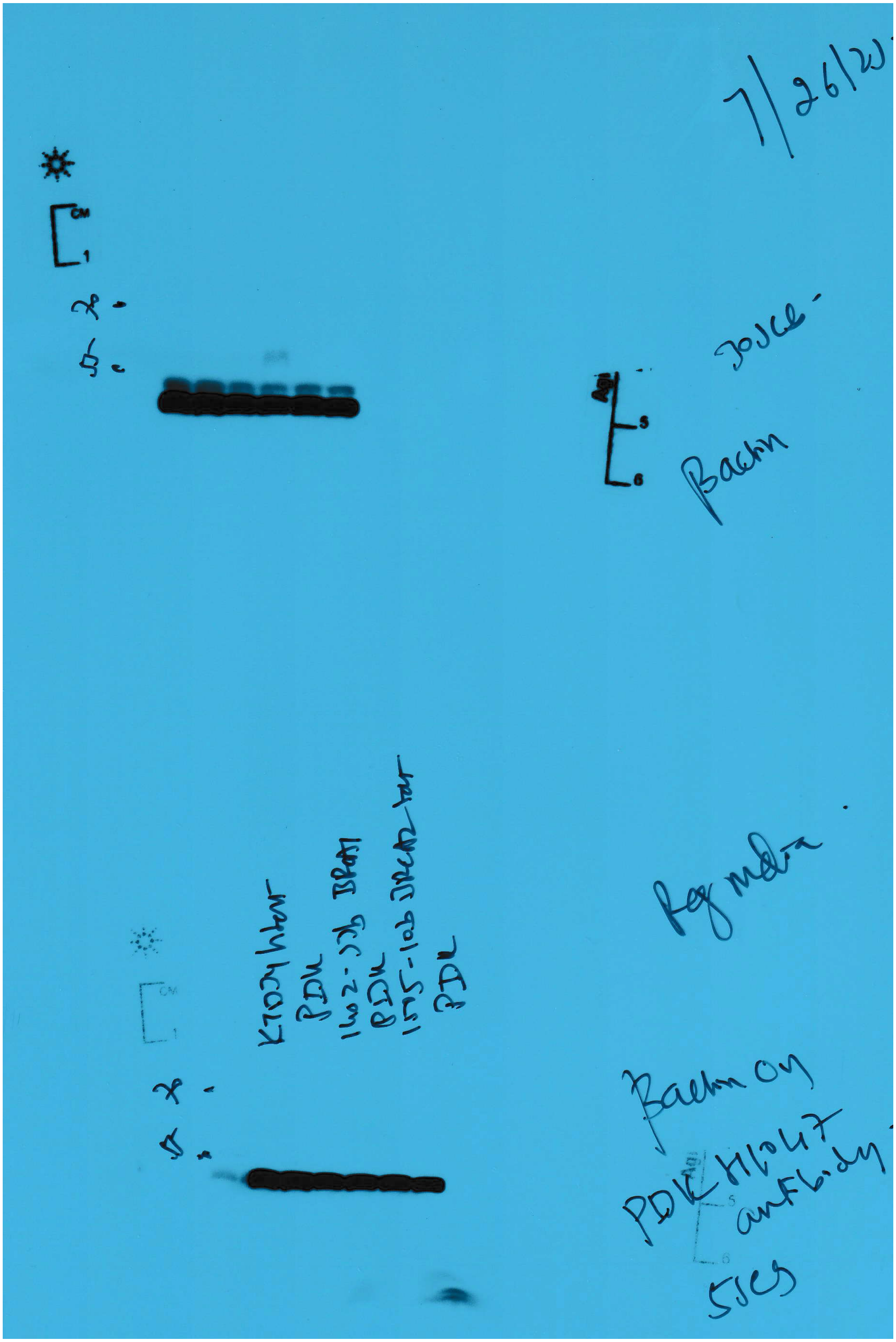
Joyce
Bactin
Reg media
Bactin on
PDK H1047
antibody

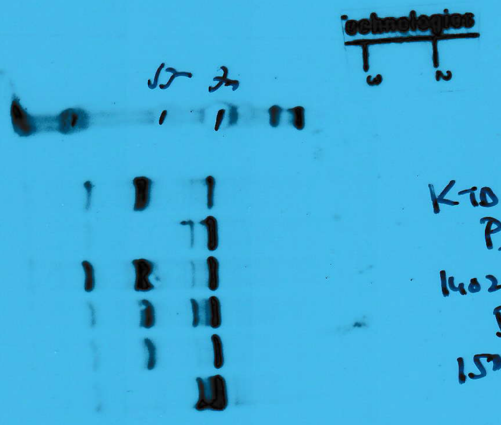


Capt-1

K7034 tert
PDK
1402-52b [illegible]
PDK
1505-6b [illegible]
PDK

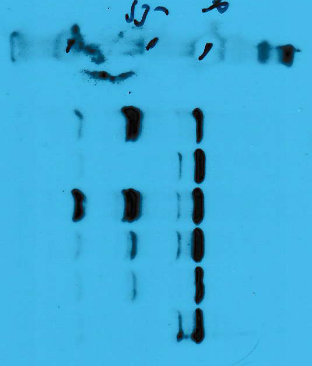

Capt-2

K7034 tert
PDK
1402-52b [illegible]
PDK
1505-6b [illegible]
PDK

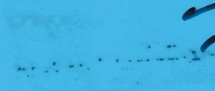

PPLO

7/6/20

2 min

SF media 0/2

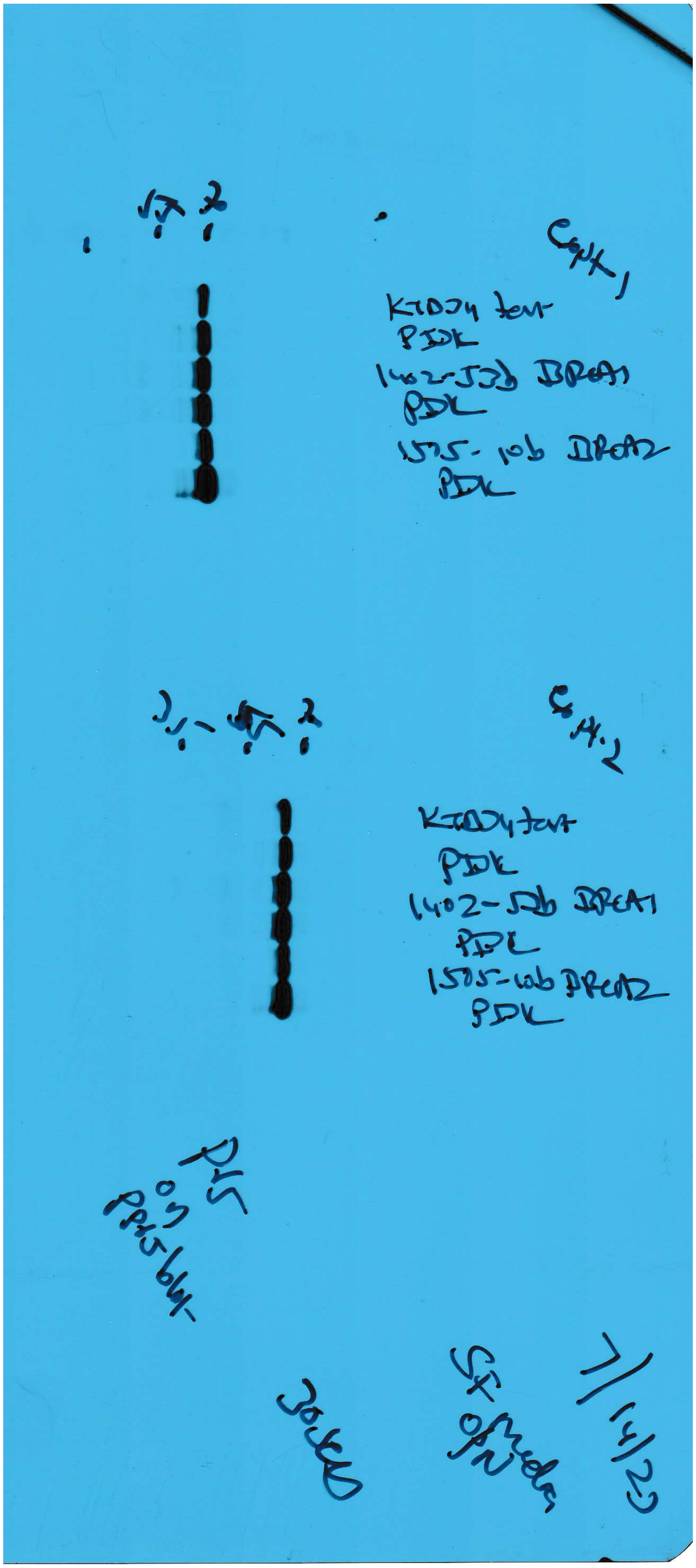

Expt 1
PDK
PDK
PDK
Expt 2
PDK
PDK
PDK

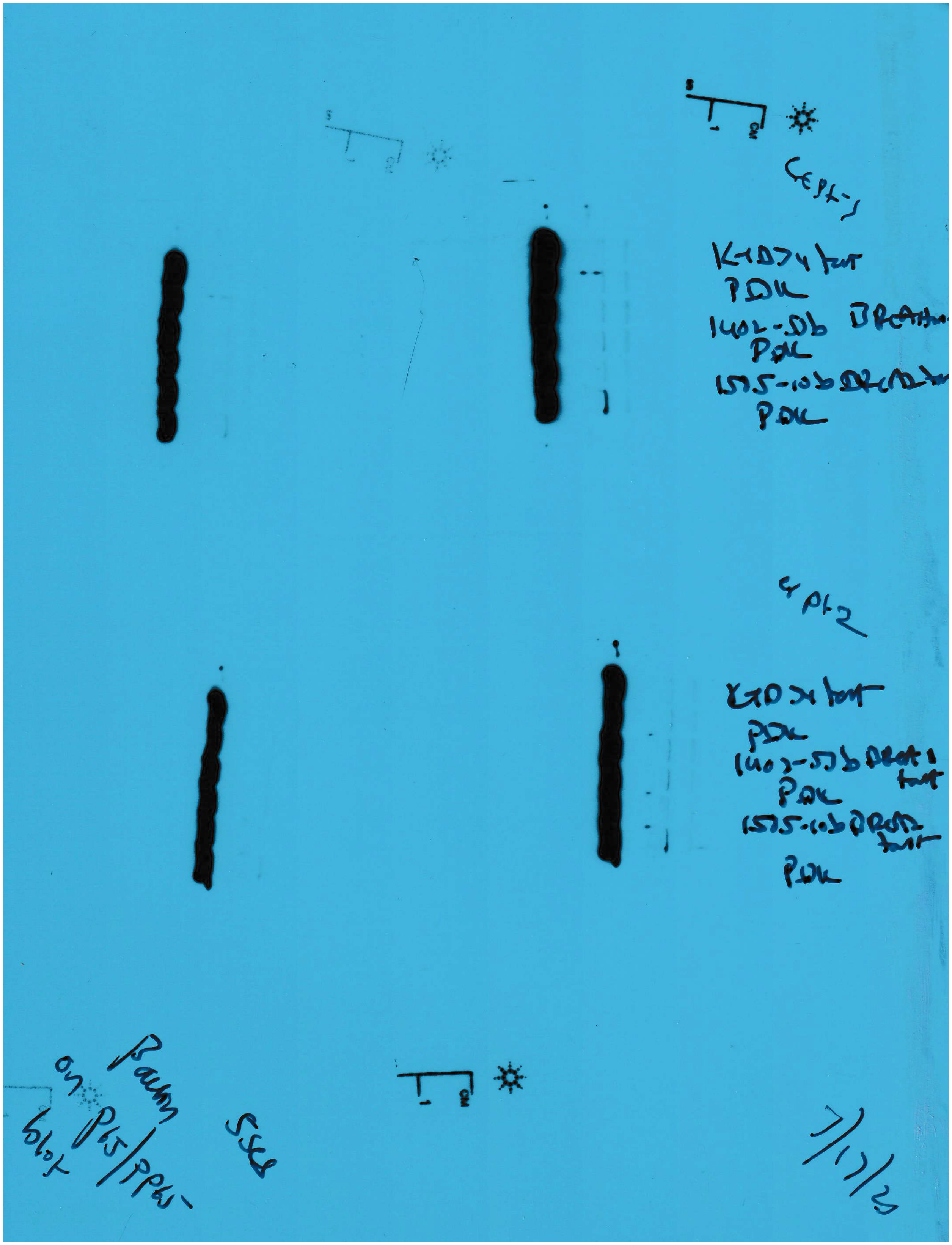
PDK
PDK
PDK
PDK
PDK
PDK
Baum
on P65/PP65
blot
